# Comparative investigation of the Diabetic foot ulcer microbiome

**DOI:** 10.1101/2021.01.21.427721

**Authors:** Amr T. M. Saeb, Hamsa T. Tayeb, Samir Ouizi, Majed S. Nassar, Assim Alfadda, Udaya Raja G. K, Balavenkatesh Mani, Amira Youssef, Mohammed S. Alsuabeyl

**Author notes:** Corresponding Author (1): Amr T. M. Saeb, Ph.D., Genetics and Biotechnology Department, Strategic Center for Diabetes Research, College of Medicine, King Saud University, KSA. Corresponding Author (2): Hamsa T.Tayeb PhD, Scientist, Department of Genetics, Research Center- Riyadh Chairperson in The Saudi Human Genome project (SHG) King Faisal Specialist Hospital & Research Center, MCD: 44117, EX: 77607/32954, P.O.Box 3354 Riyadh 11211 Saudi Arabia, MBC-03-06, or.

## Abstract

**Background:** Many factors may affect wound healing in Diabetic foot ulcer, namely, microbial density, microorganisms, microbial synergy, the host immune response, and infected tissue quality.

**Methodology:** This study used a cross-sectional design. We assessed 38 Subjects with DFUs, 23 neuroischaimic, and 15 neuropathic, for microbiota colonizing the DFU utilizing traditional cultures and 16S gene sequencing methods. All the relevant clinical factors were collected. Wound swabs were collected for both traditional microbiological analysis, direct swab (DSM), and cultured (CM) microbiome analysis. DNA isolation, and 16SrRNA hypervariable regions were amplified. Bioinformatics analysis was performed using IonReporter Software, statistical analysis, and diversity indices were computed with vegan R-package.

**Results:** The traditional microbiological method was able to detect a maximum of one or two pathogens, and, in some cases, no pathogen was detected. The total number of the observed species was 176. The number of identified species was higher in the cultured microbiome (155) than the direct swab microbiome (136). Diversity analysis indicated that biological diversity is higher in the cultured microbiome compared with the DSM. The Shannon H index was 2.75 for the cultured microbiome and 2.63 for DSM. We observed some differences in the major bacterial taxa amongst neuroischaimic and neuropathic DFU microbiomes.

**Conclusions:** Cultured microbiome is superior to both the traditional method and direct swab microbiome. The Neuroischaimic group showed higher values for the tested diversity indices than the Neuroischaimic group. Neither Cluster Analysis nor Principal Component Analysis showed apparent clustering amongst the two types of ulcers.

## 1 Background

Diabetic foot ulcers are a very important complication for diabetic patients. It is the most common cause for both hospitalization and lower extremity amputations in diabetics (1, 2). One-fourth of diabetics experience at least one diabetic foot ulcer (DFU) throughout their lifespan, and 12% of these infected upon presentation (3–5). Indeed, high wound microbial bioburden could lead to critical wound colonization that in turn lead to an overt infection that may be accountable for delaying the wound healing process and affecting the attainment of antimicrobial therapy (6). Accurate identification of causative pathogens is crucial for successful systemic antibiotic therapy of diabetic foot infection (7, 8). Moreover, it is imperative to fully identify and describe the complete collection of microorganisms colonizing the DFU to establish their role in impairing or enhancing wound healing. Many factors may affect the outcome of wound healing. These include microbial density, microorganisms, microbial synergy, the host immune response, and the quality of infected tissue (9). Furthermore, lots of investigations emphasized the polyclonal nature of diabetic foot infections (10, 11). Moreover, genotypically unrelated bacteria, which live in a symbiotic relationship in the wound, can produce a pathogenic community. Though individual bacterial species of these communities may not cause disease when they occur individually, when they congregate into a synergistic community, it can function as well-known pathogens with the ability to form a microbial biofilm that and sustain chronic infections (10). Molecular microbiological techniques have exposed the lack of precision of wound cultivation-based study findings (12). Microbial studies based on amplification and sequencing of 16S ribosomal RNA (rRNA) gene are very accurate and eliminate any biases associated with cultivation-based studies (13–15). 16S rRNA gene exists in all bacteria’s genomes and contains both variable and conserved regions that allow wide range identification and classification of bacteria through simple PCR amplification. The term microbiome refers to the collection of microbes and their genes exist in certain site while metagenomics refers to the shotgun description of total DNA, and now is progressively applied to investigations of marker genes such as the 16S rRNA gene (16). Studying the metagenomics (microbiome) of DFU allows precise evaluation of microbial load, microbial diversity, pathogenicity, and microbial community and structure within the wound (17). Thus, the metagenomics of DFU allows the understanding of the role of microbiota in DFU consequences and complications. There is no doubt that the wound sampling technique is fundamental to provide the clinicians with reliable data that can aid in taking the required steps toward infection control and wound healing. Traditional culturing approaches are very biased as a diagnostic tool toward effortlessly cultured microorganisms such as *Staphylococcus aureus* and against fastidious bacteria such as anaerobes that may have adverse effects on wound healing (10). Moreover, microbiology laboratories should realize the importance of culturing polymicrobial specimens, identifying multiple microorganisms, and selecting then testing the microorganisms for antibiotic susceptibility (9). Therefore, there is a great need for comparing traditional culture results in high-resolution molecular diagnostic methods (10). Another problem that we noticed in almost all DFU microbiome studies is the lost samples when trying to isolate DNA directly due to low DNA concentration or the low bacterial load of the swabs from the wound or the intact skin (18) (19) (15). Losing swab information is a critical matter because of losing the snapshot information in comparative studies and losing informative temporal sequential information in longitudinal studies, and it is a frequent incidence in laboratories that lack skilled technicians. There are two types of DFUs, namely, neuropathic and neuroischaemic ulcers. The neuropathic ulcers typically arise on the plantar aspects of the toes or the foot under the neuropathic foot’s metatarsal heads. The neuropathic foot is characterized by being warm, well-perfused, palpable pulses, and dry and prone to fissuring skin. While, neuroischaemic ulcers are often seen on the margins of the foot, toes’ tips, and beneath any toenails if these become overly thick in a neuroischaemic foot. The neuroischaemic foot is characterized by a lower temperature, weak or no pulse, and shiny, thin, hairless skin. Besides, it comprises atrophy of the subcutaneous tissue and painless because of Neuropathy. However, the major difference between the two types of diabetic feet is the absence or presence of ischemia that can be established by the ankle-brachial pressure index (ABPI) and doppler arterial waveform (20) (21). There are differences in the causative organisms of DFU. These differences occur due to several reasons such as geographical and environmental variations, duration of infection, or the types and severity of the infection. The majority of the studies implicated *Staphylococcus aureus* as the main causative agent of DFU infections (10). Other studies reported the prevalence of gram-negative aerobes as the major infectious pathogens (22). Lots of studies suggested a minimal role of anaerobes with Bacteroides species as an exception (23) (24), While most reported studies on diabetic foot infections have been published in developed countries, the microbiology of diabetic foot infections in Saudi Arabia has not been extensively studied. Because of that our study is aiming to compare the DFU microbial load, microbial diversity, pathogenicity detected by 16S rRNA gene sequencing to those obtained from total (swab) microbiome and cultured microbiome against conventional microbiology culturing techniques routinely used in clinical setups. In addition to Comparing between neuroischaimic and neuropathic DFU core microbiomes. Furthermore, to our knowledge, this is the first study to investigate the DFU in the Arabian Gulf region.

## 2 Materials and Methods

### Design, setting, and sample

The institutional review board of the King Saud University, College of Medicine Riyadh, Kingdom of Saudi Arabia, approved this study and was performed following the Helsinki Declaration. The participants provided written, informed consent for joining this study. Samples were collected between March 29, 2016, and December 7, 2017, in the Diabetic foot unit, University Diabetes center at King Saud University. The following criteria were used: 18 years of age or older; neuropathic or neuroischaimic DFU; free of systemic antibiotics over the past two weeks; negative for clinical signs of infection; negative for wound deterioration; and negative for osteomyelitis. All wounds were examined for the presence and severity of infection by a trained diabetologist using the PEDIS (Perfusion, Extent, Depth, Infection, and Sensation) classification system (25) (2). The following clinical information was collected at the time of sampling by clinicians: demographic data, diagnosis at admission, PEDIS grade of wound severity, topography, number, ulcer size (including surface area and depth of the wound), diabetes type, duration of Diabetes, HbA1C value, previous history of DFU, other diabetes complications, medications, and antibiotic therapy in the 14 previous days. When patients had several wounds, the sample was taken from the most severe wound. DFU management was performed according to international guidelines independently of our study results (25).

### DFU swab collection

Ulcer specimens were attained using the Levine technique and established protocols described before (19) (18). Each DFU was cleansed with non-bacteriostatic saline; three different swabs were rotated over a 1 cm^2^ area of viable non-necrotic wound tissue for 5 seconds by applying enough pressure to extract wound tissue fluid. One regular swab was directly placed into a tube containing 2 ml of sterile PBS and immediately placed on ice for DNA isolation. The second swab Amies with charcoal transport swab (Copan, Brescia, Italy), was sent to the clinical laboratory at King Abdelaziz university hospital for standard aerobic and anaerobic culture. The third swabs were vortexed in 1 mL tryptic soy broth and then diluted and plated on Columbia blood agar, eosin-methylene blue agar, CHROMagar MRSA, and Brucella Agar supplemented with blood, hemin, and vitamin K. All organisms that grew in both aerobic and anaerobic cultures were collected and suspended 2 ml of sterile PBS and stored for DNA isolation (**Supplementary Figure 1**).

### 16S rRNA gene sequencing, quality control, and analysis

Bacterial cells were released in the 2 ml of PBS were collected by centrifugation, resuspended a 0.5 mL ReadyLyse lysozyme solution (Epicentre), and then sample incubated for one hour at 37°C with shaking. Samples were then processed in a TissueLyser (Qiagen) at maximum speed for 2 min, followed by a 30- min incubation at 65°C with shaking. DNA was extracted from samples using Maxwell® 16 automated system (Maxwell® 16 Cell DNA kits, Promega, Madison, WI, USA). Extracted DNA samples were quantified using NanoDropTM 2000 UV-Vis Spectrophotometer (Thermo Fisher Scientific, Waltham, MA). The quality of extracted DNA samples was also checked by loading one μl of DNA in 1% Agarose Gel (Bio-Rad, California, USA). The extracts were stored at -20 C until use.

### 16SrRNA gene hypervariable regions amplification

Library preparation using 50ng of extracted DNA samples involved the amplification of the 16S region using 16S Ion Metagenomics Kit ™ (Thermo Fisher Scientific, Waltham, MA). Amplification comprised of two different polymerase chain reactions targeting V2-4-8 and V3-6, 7-9 hypervariable region of 16S rRNA following the manufacturer’s instructions performed using Veriti™ 96-Well Thermal Cycler (Thermo Fisher Scientific, Waltham, MA). A 2 % Agarose gel (Bio-Rad, California, USA) was prepared, and five μl of each amplified product was run along with manufacturer-provided positive control, negative control, and 50 bp markers (50 bp step ladder, Promega) to confirm the respective expected product sizes. PCR cleanup was performed for all the amplified products to remove non-specific products, excess nucleotides, and primers using Wizard^®^ SV Gel and PCR Clean-Up System (Promega, Madison, WI, USA). Cleaned amplified products were again run (5μl loaded) on 2% Agarose gel (Bio-Rad, California, USA) to confirm no further contamination. Equal volumes of cleaned V2-4-8 and V3-6, 7-9 amplicons of each sample were pooled together. The PCR amplifications were performed using Applied Biosystems VeritiTM 96-Well Thermal Cycler under the following cycling conditions: 95°C for 10 min, then Cycle 18–25 cycles 95°C for 30 sec, 58°C for 30 sec, and 72°C for 20 sec. Followed by a final extension step at 72°C for 7 min, then Hold at four °C overnight. The PCR amplified products were used for library preparation using Ion plus Fragment Library kit with sample indexing using the IonXpress Barcode adapters 1-16 kit following the manufacturer’s instructions. The Library concentration was measured using the Ion universal library quantification kit and diluted to 13 pM. The Diluted libraries were equally pooled together and used as a template for emulsion PCR. Emulsion PCR and enrichment of template-positive particles were performed using an Ion PGM Hi-Q OT2 Kit (Thermo Fisher Scientific) and the Ion OneTouch 2 system (Life Technologies) according to the manufacturer’s instructions. The enriched template-positive Ion sphere particles were loaded onto an Ion 318 chip v2 (Thermo Fisher Scientific) and sequenced on an Ion PGM (Thermo Fisher Scientific) using an Ion PGM Hi-Q Sequencing Kit (Thermo Fisher Scientific) according to the manufacturer’s instructions to ensure a high degree of sensitivity required and the deal with the high complexity of our samples under investigation.

### DNA sequence analysis

Data analysis was carried out with automated streamlined software’s Torrent Suite Software v5.4 (TSS) and IonReporter Software v5.6. Torrent Suite Software performed sequencing base call from IonTorrent PGM instrument and IonReporter for annotation and taxonomical assignments. Standard analysis parameter used in Torrent Suite Software at primary base calling analysis including trimming low-quality 3’ ends of reads, removing duplicated reads, filtering out entire reads with an average quality score less than Q20 (base call quality), removing of adapter sequence, removal of lower-quality 3’ ends with Low-Quality Scores, removing short reads, removing adapter dimers, removing polyclonal reads. Q20 is the Phred scale score based on error probability (-10×log10). Q20 corresponds to a predicted error rate of one percent. AQ20 is the read length at which the error rate is 1% or less. IonTorrent PGM generated sequences saved in binary alignment map (uBam) file format consist of base call sequences, flow (flow gram), and qual (quality score). IonReporter Uploader plugin used to transfer all samples of high-quality data from Torrent Suite Software to cloud Ion Reporter Software.

Two groups of samples Group-A & Group-B, 38 samples in each group analyses launched in IonReporter Software with 16S Metagenomics Analysis Workflow. Metagenomics workflow contains curated Greengenes and premium curated MicroSEQ ID 16S rRNA reference databases. Greengenes databases manually custom-curated from the available existing public library because it is relatively large references containing more than 1.2 million references. The Greengenes database is mostly the aim towards taxonomy information for family level, and above, taxonomy information for genus and species levels was missing for almost 1 million references. A custom utility was used to curate Greengenes for genus and species-level information by querying NCBI. The final library contained 215,967 references directly from Greengenes, and 146,822 references with updated genus and species names from NCBI, totaling 362,789 references. Both the databases library was converted into a BLAST compatible format for the analysis.

The analysis pipeline includes annotation, taxonomic assignment, and reads classification through alignment to either the curated MicroSEQ ID or curated Greengenes databases. IonReporter identifies chimeric sequences based on a built-in algorithm and removes them before passing them to subsequent downstream analysis. Each group samples analyzed together, and reads were classified at the species level with the taxonomical order with 16S Metagenomics Analysis Workflow. Analysis steps include trimming primer and length check after trimmed reads, processed reads placed in a hash table. This hash table contains all the unique reads and their abundance or copy number. When adding a new read to the hash table, the read will have a copy number of 1 if it does not exist in the table already, and if it does, the copy number will be increased by 1. After adding all reads, the hash table contained all unique reads, each with a copy number. Followed by all unique reads are converted to FASTA file format. Each read has a unique ID to keep analysis track of results throughout the process. A multistage BLAST search of reads against databases, this algorithm generates the input files and starts one BLAST process for each sample. BLAST parameters included E-value equal to 0.01; the result E value is higher than 0.01 was omitted in the results output. These final ion-reporter-generated results consist of species-level OUTs (operational taxonomic unit) identification, which also provides primer information, classification information, percent ID, and mapping information (26, 27). All sequence data were submitted to NCBI with an SRA accession: SUB4604842 and BioSample accessions numbers from SAMN10174890 to SAMN10174927 under the BioProject PRJNA494939

### Statistical and phylogenetic analysis

Statistical analysis was performed as described in (Saeb et al., 2019) (26). Data were analyzed using Statistical Package for Social Studies (SPSS 22; IBM Corp., New York, NY, USA) and Past 3.21 software. The demographic data, frequency, and means were computed for each patient and group. The clinical parameters were averaged for each patient and the three glycemic groups. Comparisons amongst groups were evaluated by Chi-square (for categorical data), Mann-Whitney (for pairs of groups). The continuous variables were expressed as mean ± standard deviation. The t-test was used for continuous variables with a normal distribution, and the Mann-Whitney test was used for non-normal distribution.

Shapiro-Wilk test was used to assess the normality of the data. A p-value <0.05 was considered statistically significant. All diversity indices were calculated using statistical software R, vegan R-package, and Past321 software (28). Cluster analysis (Unweighted pair-group average UPGMA) using Bary-Curtis, Manhattan, Elucidation, and correlation similarity indices with bootstrap number=1000 were performed using Past321 software. The principal component analysis using variance-covariance and correlation matrix was also performed using Past321 software.

## 3 Results

### 3.1 Recruitment of the study subjects

Only 38 subjects fulfilled the study protocol, after excluding any subjects who had taken antibiotics in three months. The final study cohort included 26 subjects with neuroischaimic DFU and 12 subjects with neuropathic DFU. The age ranges for the selected study cohort were 41–89 years. Males were more predominant than females, 29 and 9 respectively. The mean diabetes duration was 19 years, ranging from 1 to 36 years. All patients suffered from diabetes-associated complications. In addition to Neuropathy, all patients suffered from a range of additional complications that ranged from 1-6 complications. HbA1c levels ranged between 5% and 18.8%, with a mean of 9.8%. The wound surface area ranged between 0.25-56 cm^2^ with a mean of 13.79 cm^2^. The wound depth ranged between being superficial wound to be deep to the bone. The cause of the wound different from reasons such as footwear or trauma. The ankle-brachial pressure index (ABPI) ranged from 0.23 to 1.76, with a mean of 0.76. All medications taken during the study period are listed in the **S1 table**.

### 3.2 Sequencing data and taxonomic assignment of the sequence reads

The bacterial composition of the DFU samples of subjects in the current study was determined by 16S rRNA gene amplicon analysis using an Ion Torrent proton machine. After applying the read quality and length filters, 478470 raw reads were obtained for the cultured microbiome. The number of reads per sample was 728–42323, with a mean of 12591 reads per sample. After applying the read quality and length filters, 337657 raw reads were obtained for the direct swab microbiome. The number of reads per sample was 1279 –42323, with a mean of 8886 reads per sample. The IonReporter software automatically removed the chimeric reads; further, UCHIME, Chimera Slayer, and Decipher programs identified 0.5% chimeric reads later excluded. The average read length was 170 base-pairs. The sequences were assigned to 176 operational taxonomic units (OTUs), using a cutoff distance of 0.05, by sequence comparison to the updated databases (**Figure 1**).

**Figure 1.**
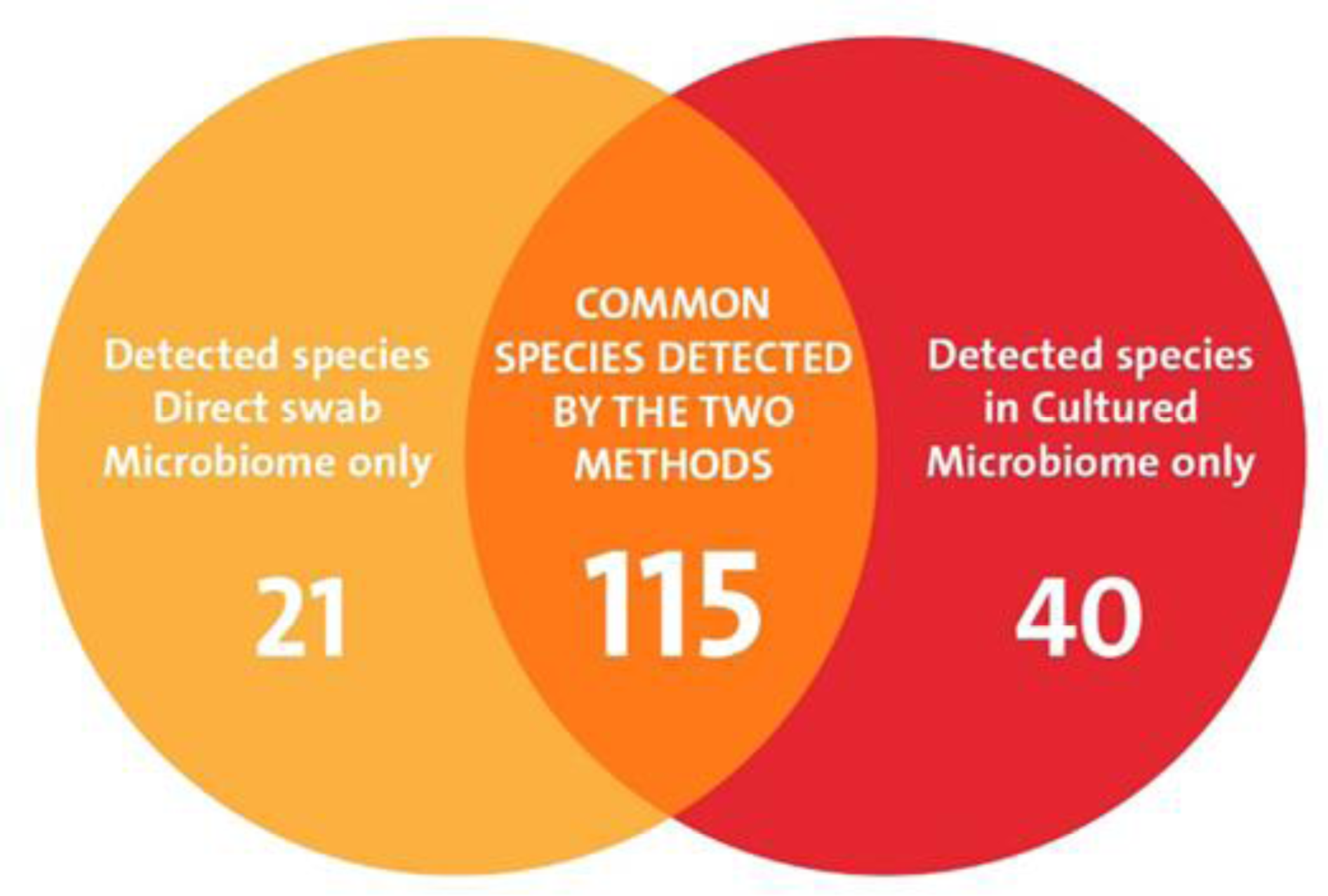
Distribution of the observed species (OTUs) in the two DFU microbiome methods.

### 3.3 The core DFU microbiome: class level

In the sampled population (38 individuals), three classes, all from the domain Bacteria, represented the core taxa (95% of reads) in the DFU microbiome. These were Bacilli, Actinobacteria, and Gammaproteobacteria (**Figure 2a**). The class Bacilli represented 70 and 79% of the core microbiome for both methods, followed by Gammaproteobacteria 20 and 15% in cultured and direct swab microbiome methods, respectively. Actinobacteria represented 10 and 6 % for cultured and direct swab microbiome methods, respectively.

**Figure 2.**
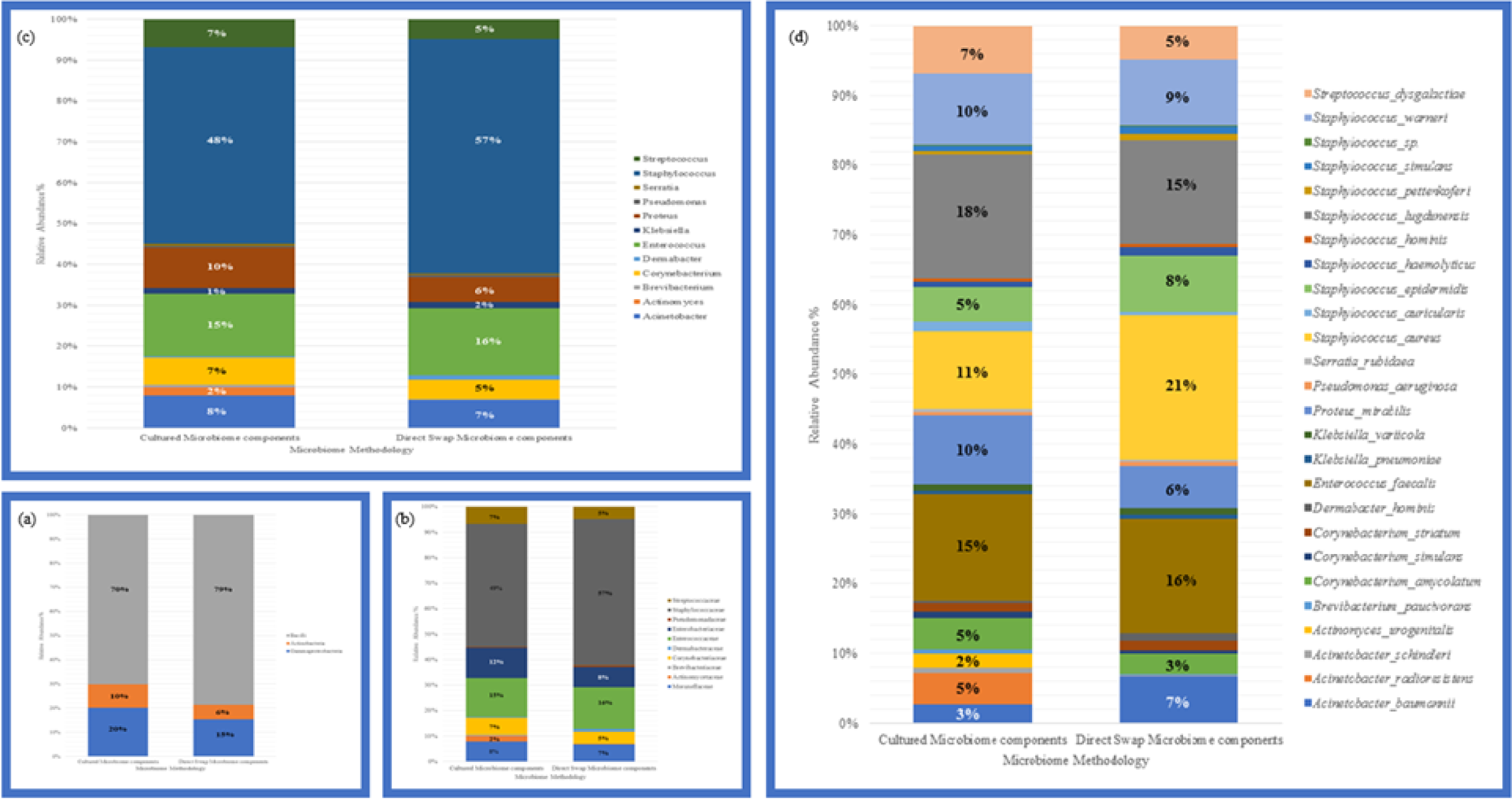
Taxonomic comparison between the core DFU microbiome for CM and DSM methods

### 3.4 The core DFU microbiome: family level

In the sampled population (38 individuals), ten families represented the core taxa (95% of reads) in the DFU microbiome. These were Moraxellaceae, Actinomycetaceae, Brevibacteriaceae, Corynebacteriaceae, Dermabacteraceae, Enterococcaceae, Enterobacteriaceae, Pseudomonadaceae, Staphylococcaceae, and Streptococcaceae (**Figure 2b**). The family Staphylococcaceae was the most abundant component of the DFU core microbiome in both methods, 49% and 57% for cultured microbiome and direct microbiome methods, respectively. The family Pseudomonadaceae was the least abundant component of the DFU core microbiome (0.38%) of the cultured microbiome method. While the family Actinomycetaceae was the least abundant component of the DFU core microbiome (0.1%) of the direct swab microbiome method.

### 3.5 The core DFU microbiome: genus level

In the sampled population (38 individuals), 12 genera represented the core taxa (95% of reads) in the DFU microbiome using the two methods. These were Acinetobacter, Actinomyces, Brevibacterium, Corynebacterium, Dermabacter, Enterococcus, Klebsiella, Proteus, Pseudomonas, Serratia, Staphylococcus, and Streptococcus (**Figure 2c**). The genus Staphylococcaceae was the most abundant component of the DFU core microbiome in both methods, 48% and 57% for cultured microbiome and direct microbiome methods, respectively. The genus Pseudomonas was the least abundant component of the DFU core microbiome (0.38%) of the cultured microbiome method. While the genus Actinomyces was the least abundant component of the DFU core microbiome (0.1%) of the direct swab microbiome method.

### 3.6 The core DFU microbiome: species level

In the sampled population (38 individuals), 26 species represented the core taxa (95% of reads) in the DFU microbiome in both methods. These were, *Acinetobacter baumannii, Acinetobacter radioresistens, Acinetobacter schindleri, Actinomyces urogenitalis, Brevibacterium paucivorans, Corynebacterium amycolatum, Corynebacterium simulans, Corynebacterium striatum, Dermabacter hominis, Enterococcus faecalis, Klebsiella pneumoniae, Klebsiella variicola, Proteus mirabilis, Pseudomonas aeruginosa, Serratia rubidaea, Staphylococcus aureus, Staphylococcus auricularis, Staphylococcus epidermidis, Staphylococcus haemolyticus, Staphylococcus hominis, Staphylococcus lugdunensis, Staphylococcus pettenkoferi, Staphylococcus simulans, Staphylococcus sp., Staphylococcus warneri and Streptococcus dysgalactiae* (**Figure 2d**). *Staphylococcus lugdunensis* was the most abundant species in the cultured DFU core microbiome method (18%). While *Staphylococcus aureus* was the most abundant species in the direct swab DFU core microbiome method (21%). The unresolved OUT that belongs to *Staphylococcus sp.* was the least abundant component of the DFU cultured core microbiome (0.16%). While *Acinetobacter schindleri* was the least abundant component of the DFU core microbiome (0.014%) of the direct swab microbiome method.

### 3.7 Biological diversity of DFU using the two microbiome methods

An overview of biological alfa-diversity and diversity indices calculated for total cultured microbiome and direct swab microbiome methods is presented in **Table 1**. As stated above, the number of identified OTUs (species) was higher in the cultured microbiome (155) compared with the direct swab microbiome (136).

**Table 1.** Diversity indices for the total Cultured Microbiome and Direct Swap Microbiome methods.

|  | **Cultured Microbiome** | **Direct Swap Microbiome** |
| --- | --- | --- |
| **Taxa_S** | 155 | 136 |
| **Dominance_D** | 0.09516 | 0.1062 |
| **Simpson_1-D** | 0.9048 | 0.8938 |
| **Shannon_H** | 2.75 | 2.684 |
| **Evenness_e^H/S** | 0.1009 | 0.1077 |
| **Brillouin** | 2.749 | 2.683 |
| **Menhinick** | 0.2241 | 0.234 |
| **Margalef** | 11.78 | 10.61 |
| **Equitability_J** | 0.5453 | 0.5464 |
| **Fisher_alpha** | 14.94 | 13.42 |
| **Berger-Parker** | 0.1735 | 0.1949 |
| **Chao-1** | 155 | 136 |

The most well-known and accepted diversity indices, i.e., Chao-1, Margalef, and Fisher alpha, indicated that biological diversity is higher in the cultured microbiome than the direct swab microbiome. For example, the Chao-1 index was 155 for the cultured microbiome and 136 for the direct swab microbiome. Moreover, The Shannon H index was 2.75 for the cultured microbiome and 2.63 for the direct swab microbiome. The Margalef index was 11.78 for the cultured microbiome and 10.61 for the direct swab microbiome. The Fisher alpha index was 14.94 for cultured microbiome and 13.42 for the direct swab microbiome (**Figures 3**). **However, only the taxa, Chao-1 index, and Margalef index showed a significant difference between the two biological diversity methods (Table 2).**

**Figure 3.**
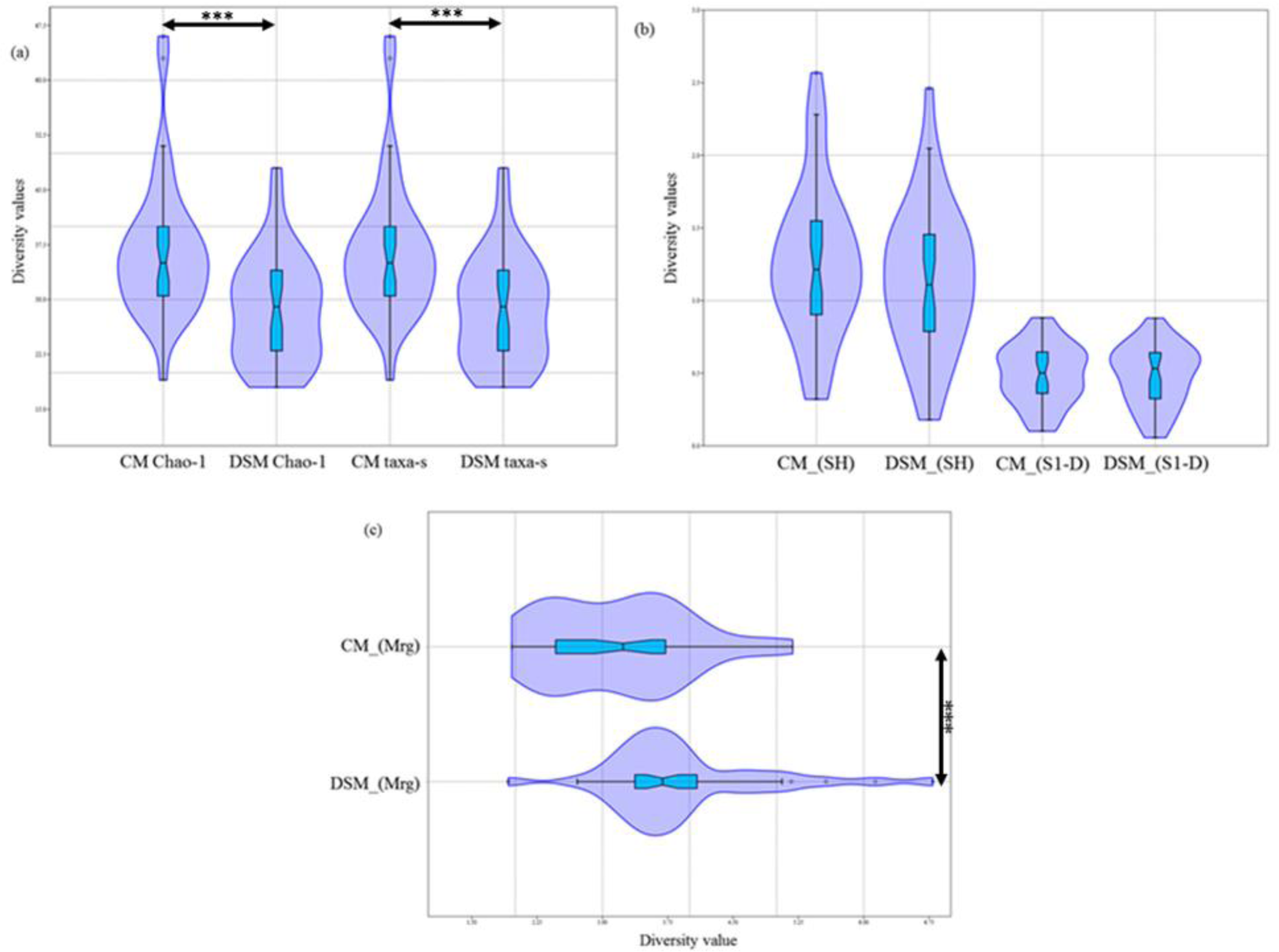
Violin plots of the diversity indices (a) Chao-1 and Taxa-s (b) Shannon H and Simpson 1-D (c) Margalef for CM and DSM methods.

**Table 2.**
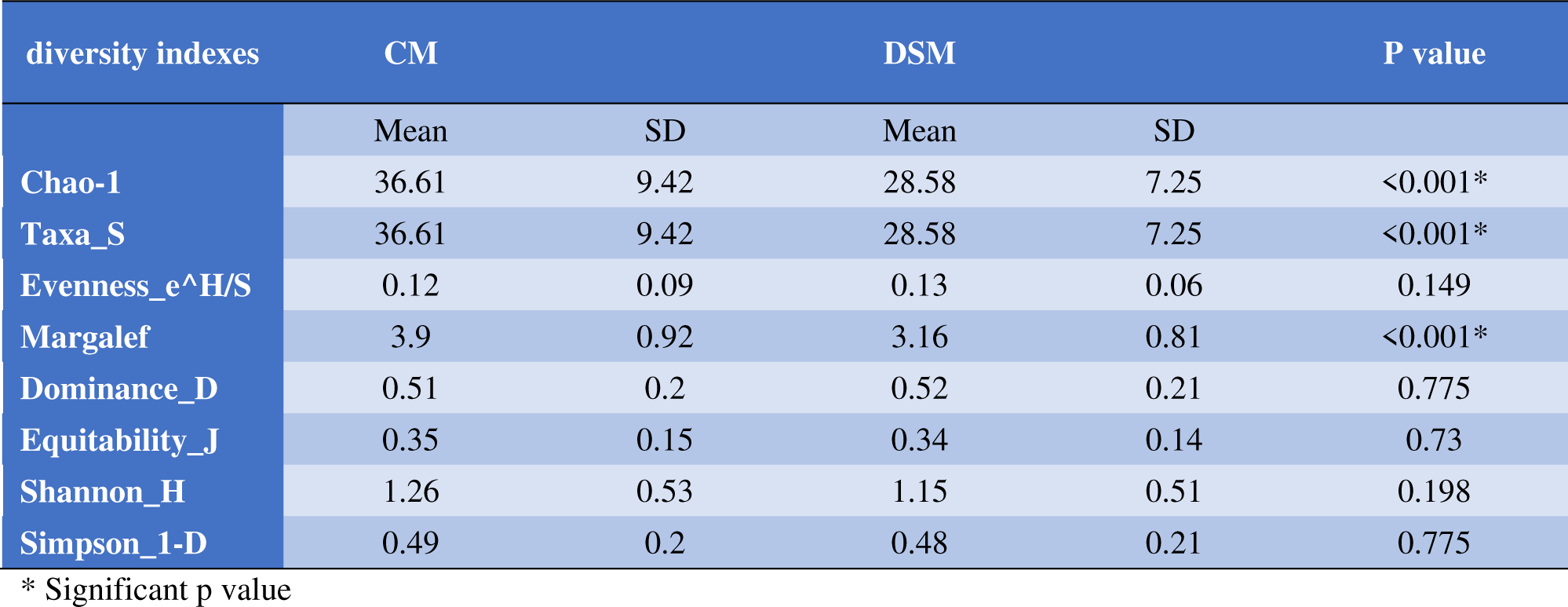
Compression between CM and DSM by different diversity indexes

| diversity indexes | CM |  | DSM |  | P value |
| --- | --- | --- | --- | --- | --- |
|  | Mean | SD | Mean | SD |  |
| Chao-1 | 36.61 | 9.42 | 28.58 | 7.25 | <0.001* |
| Taxa_S | 36.61 | 9.42 | 28.58 | 7.25 | <0.001* |
| Evenness_e^H/S | 0.12 | 0.09 | 0.13 | 0.06 | 0.149 |
| Margalef | 3.9 | 0.92 | 3.16 | 0.81 | <0.001* |
| Dominance_D | 0.51 | 0.2 | 0.52 | 0.21 | 0.775 |
| Equitability_J | 0.35 | 0.15 | 0.34 | 0.14 | 0.73 |
| Shannon_H | 1.26 | 0.53 | 1.15 | 0.51 | 0.198 |
| Simpson_1-D | 0.49 | 0.2 | 0.48 | 0.21 | 0.775 |

\* Significant p value

### 3.8 Comparison between traditional method, CM and DSM in microbe detection

**Table 3** shows a comparison between different diagnostic methods, namely, traditional microbiological method, CM, and DSM in DFU wound pathogens detection. The traditional microbiological method failed to compete with either tested microbiome method in reflecting the real microbial situation inside the DFU in all 38 cases. The traditional microbiological method was able to detect a maximum of 1 or 2 pathogens, and in some cases, no pathogen was detected. *Staphylococcus aureus* was the most detected pathogen in most cases in combination with unclassified Gram-positive bacteria. In some cases, *Proteus mirabilis, Enterococcus faecalis, Acinetobacter baumannii, Achromobacter xylosoxidans, Pseudomonas aeruginosa, Streptococcus agalactiae, Citrobacter freundii, Citrobacter koseri,* and *Serratia liquefaciens* were detected as primary or secondary pathogens. The cultured microbiome method was able to detect more wound pathogens in most of the cases (55%), while the direst swab microbiome methods detected more pathogens in (29%) cases, and they were the same in (16%) cases. The highest number of detected pathogens using the CM method (20) was detected in patient number 38, while the lowest number (2) was detected in patients 3, 14, and 21. The highest number of detected pathogens using the DSM method (13) was detected in cases number 34, 37, and 38, while the lowest number (1) was detected in patients 28 and 36. On the contrary, with the traditional microbiology method, both CM and DSM presented information about the percentages of each OUT in the core microbiome for the 38 DFU cases. For the CM method, *Staphylococcus aureus* represented the highest fraction of the core DFU microbiome in only 4 cases (10.5%), while it represented the highest fraction of the core DFU microbiome in 11 cases (29%) using the DSM method. For the CM method, *Proteus mirabilis* represented the highest fraction of the core DFU microbiome in only 5 cases (13%), while it represented the highest fraction of the core DFU microbiome in 2 cases (5%) using the DSM method. *Enterococcus faecalis* represented the highest fraction of the core DFU microbiome in only 5 cases (13%) in the CM method, while it represented the highest fraction of the core DFU microbiome in 4 cases (10.5%) using the DSM method.

**Table 3.**
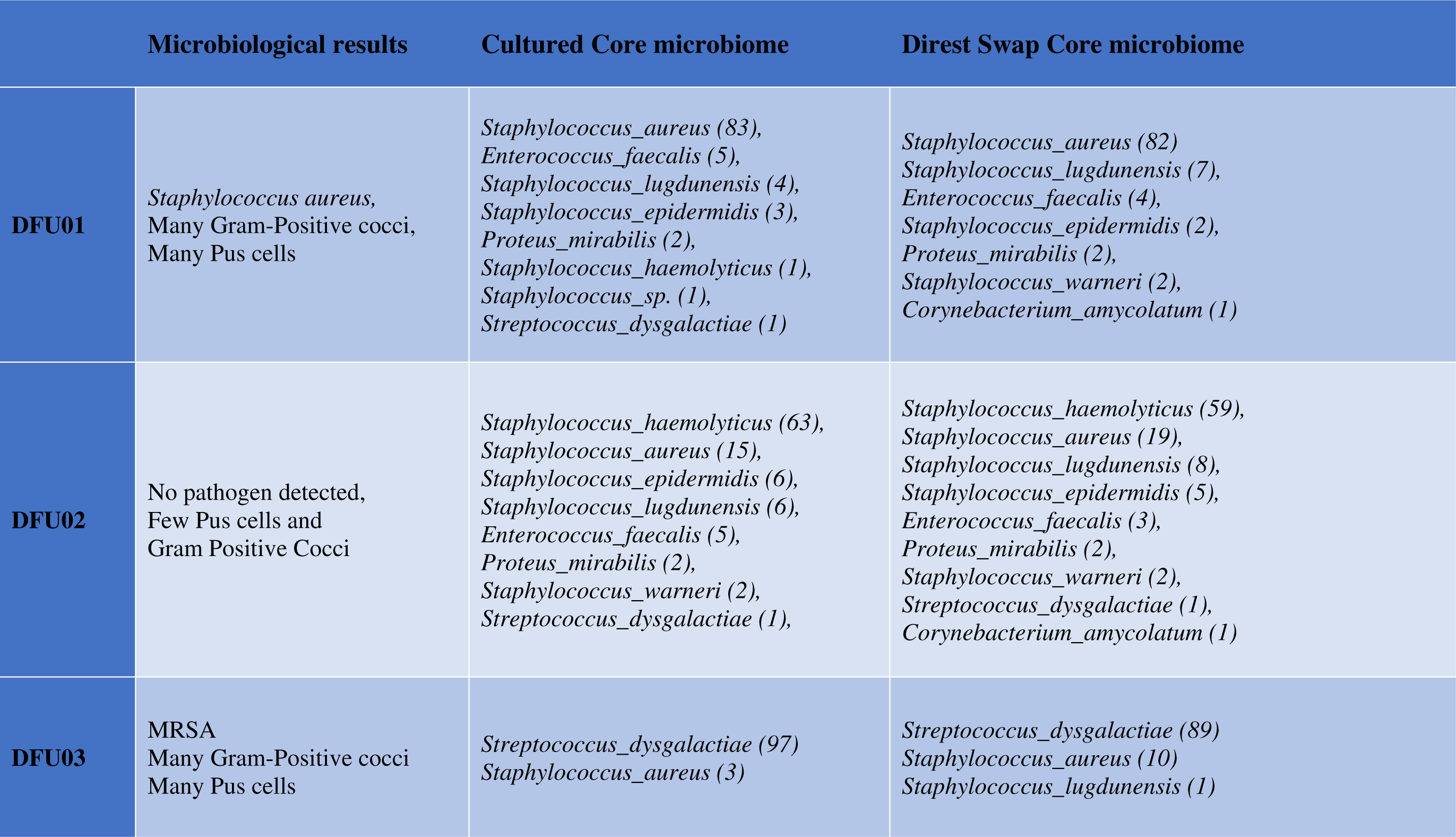
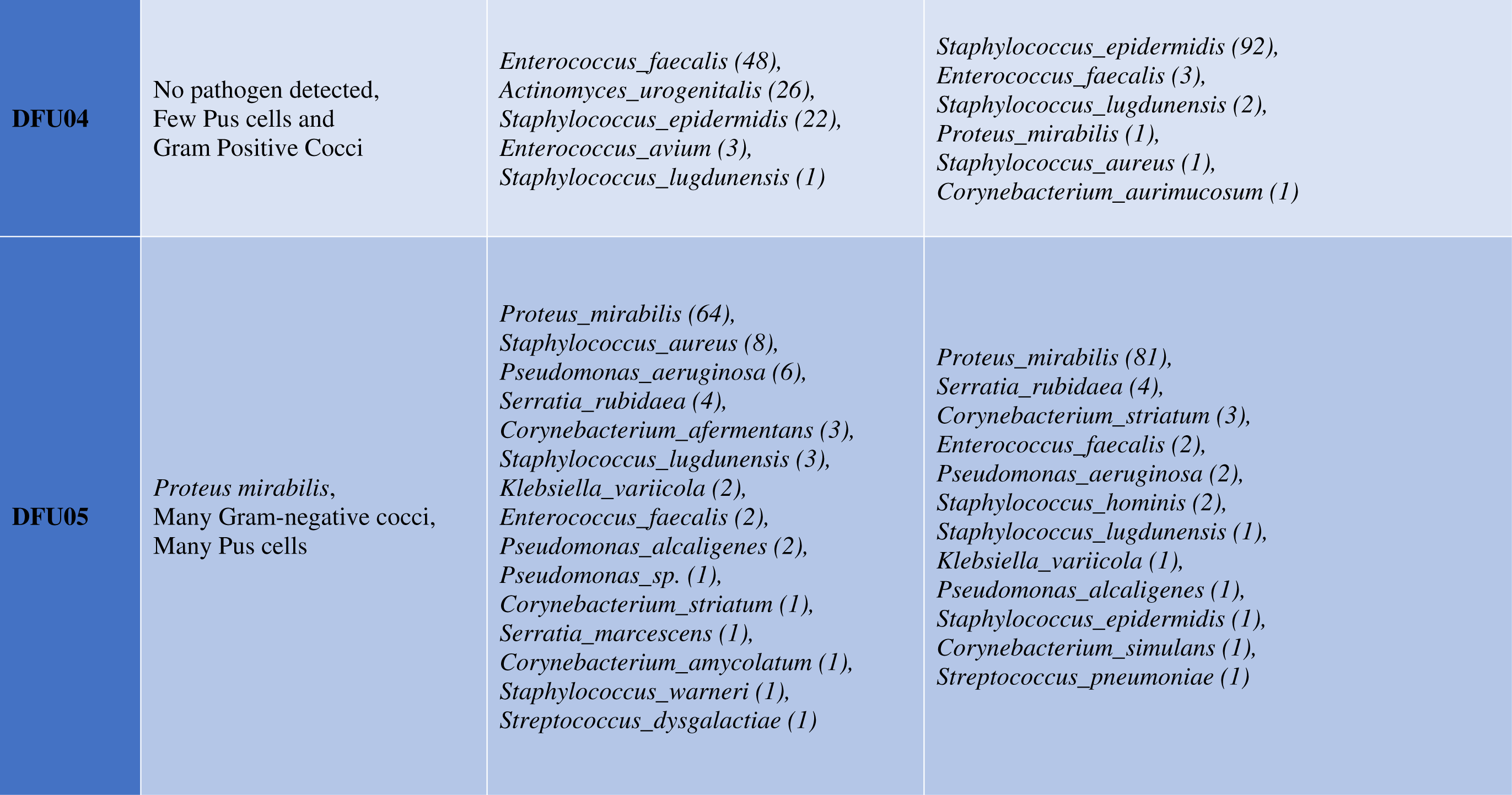
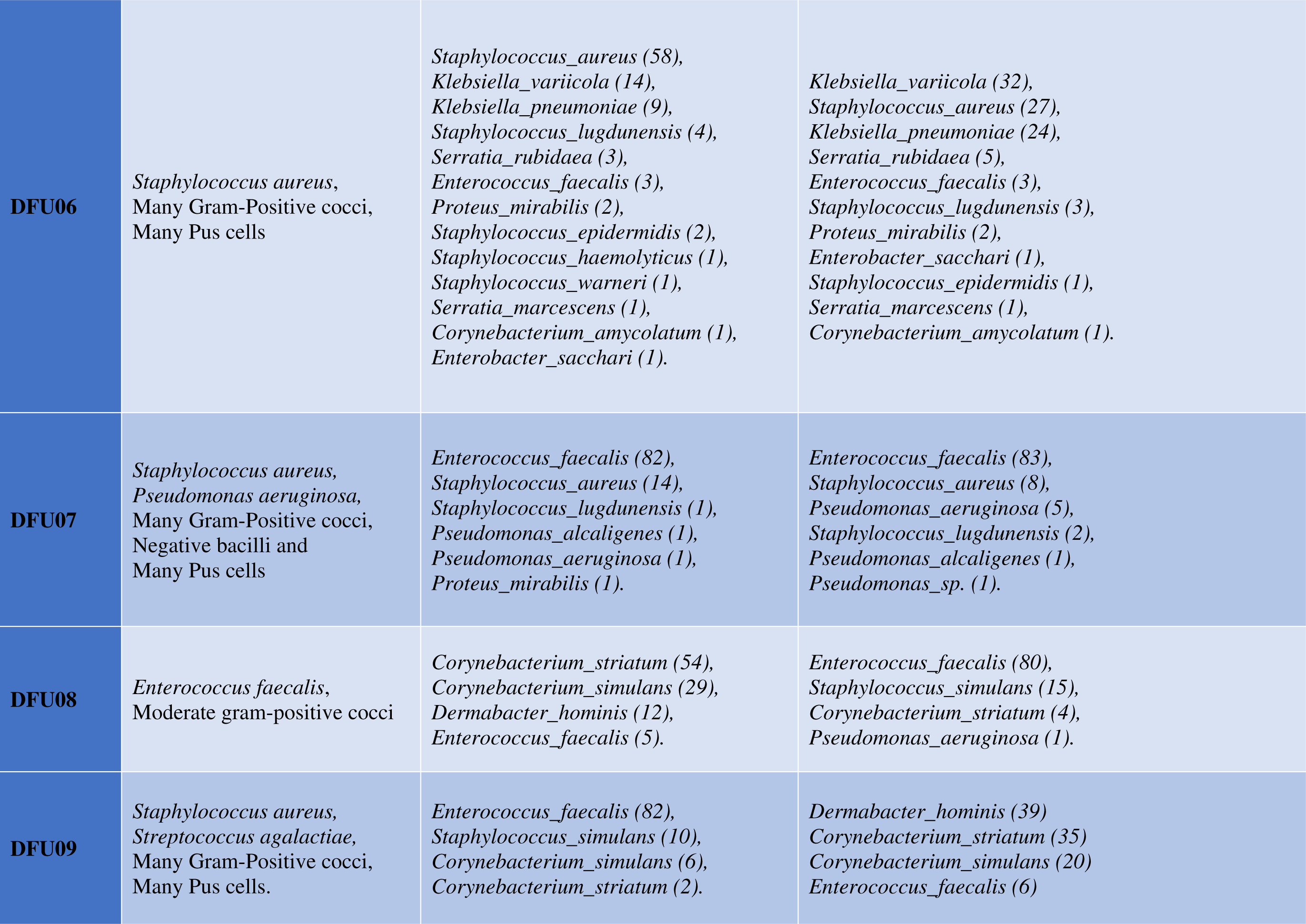
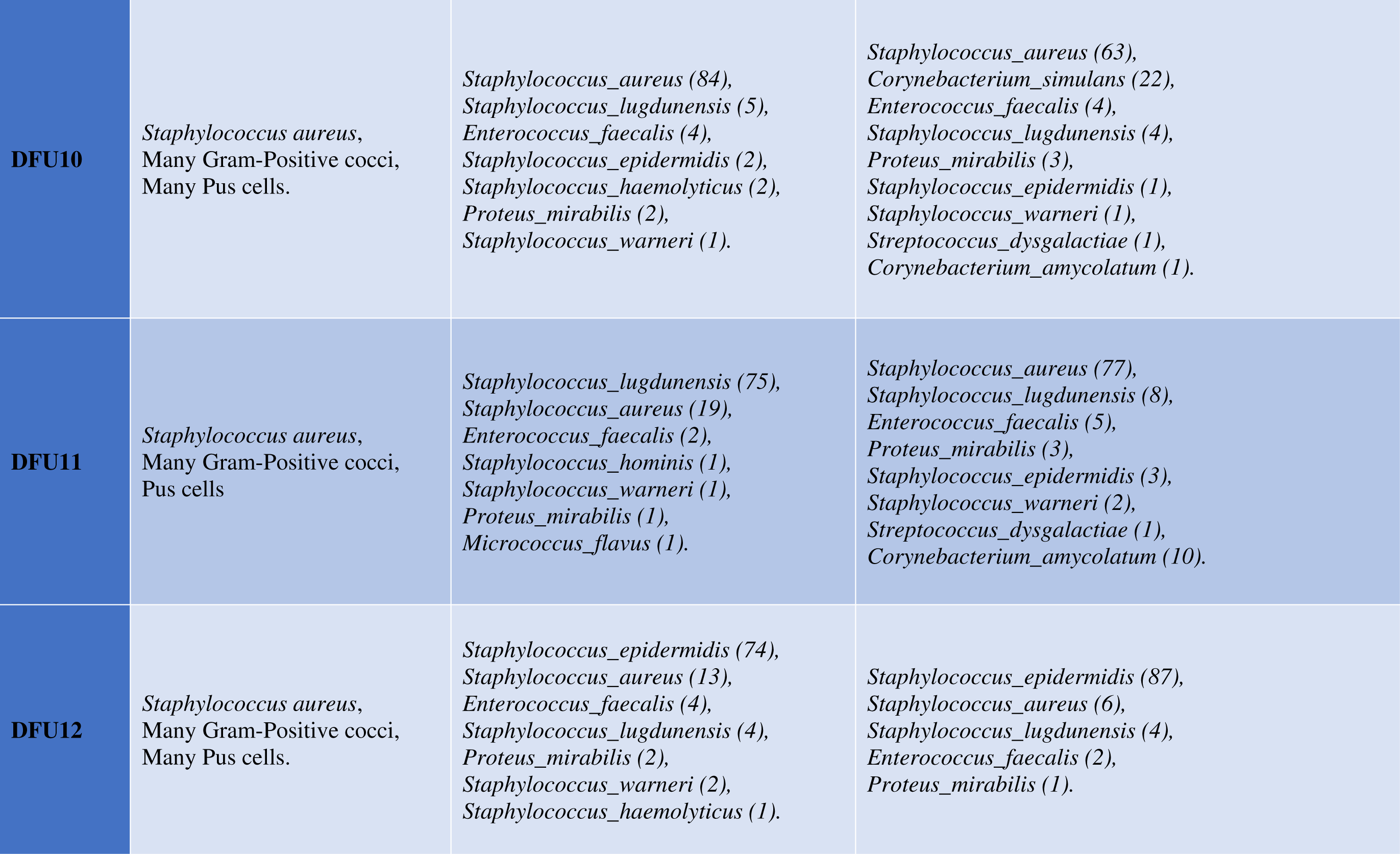
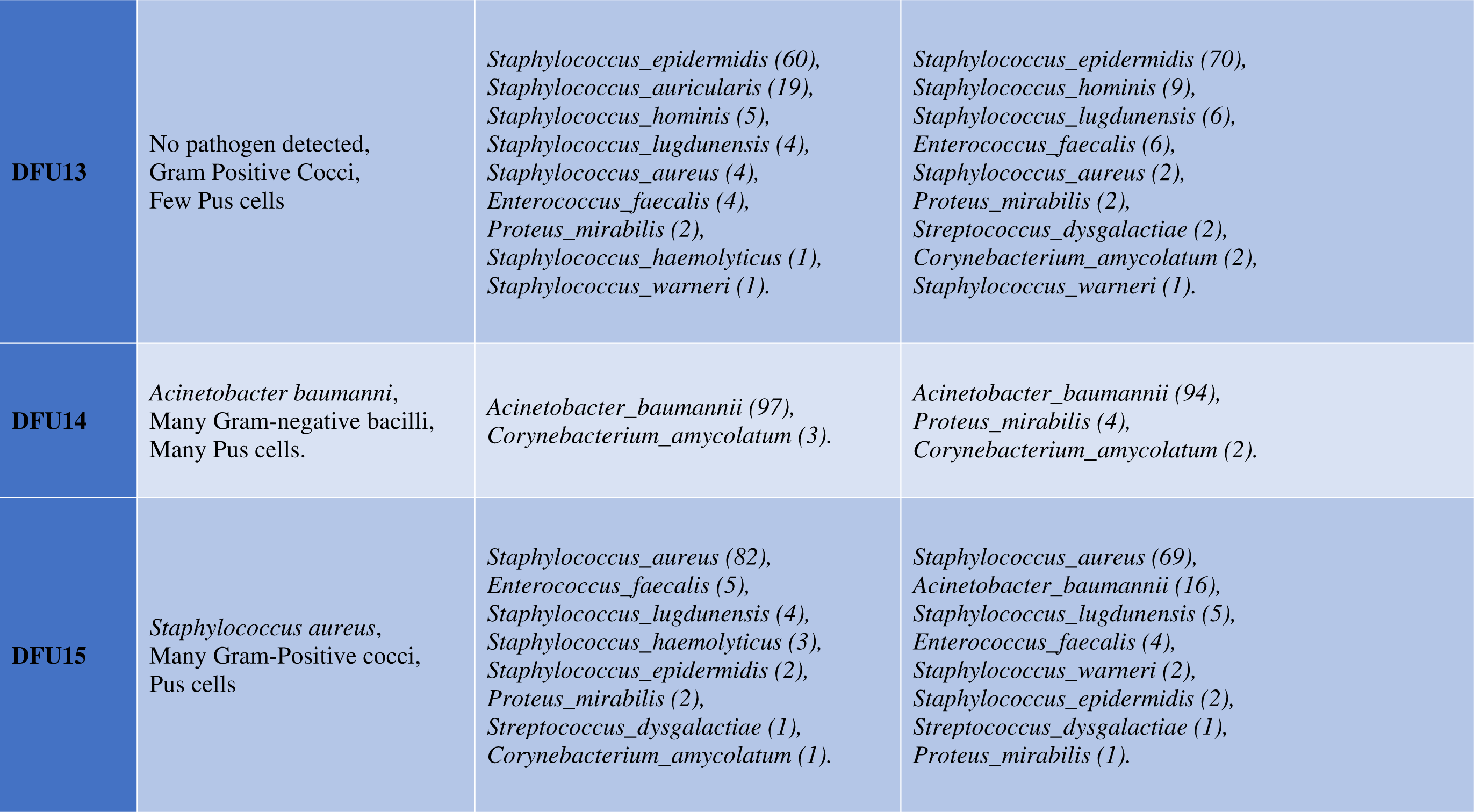
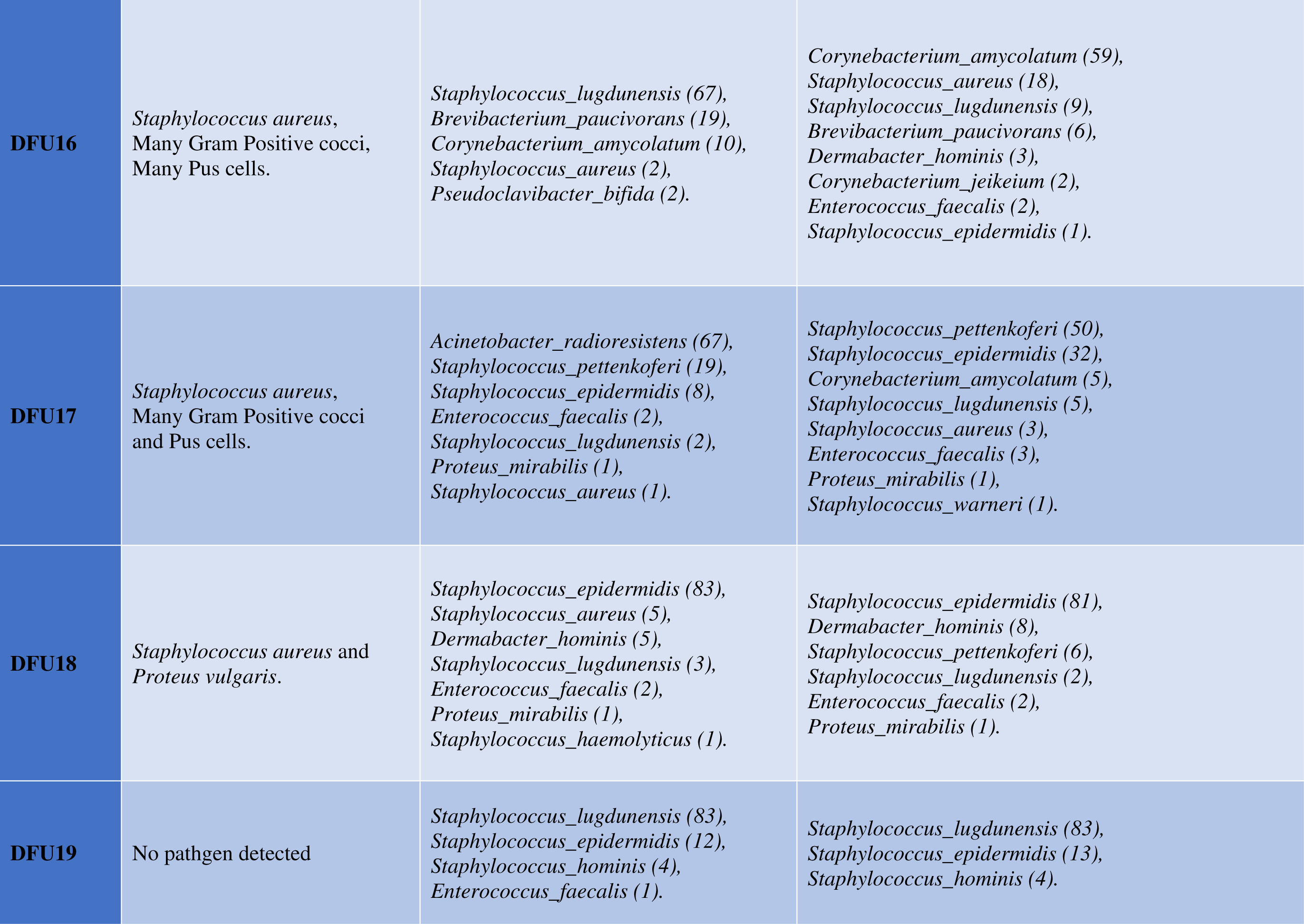
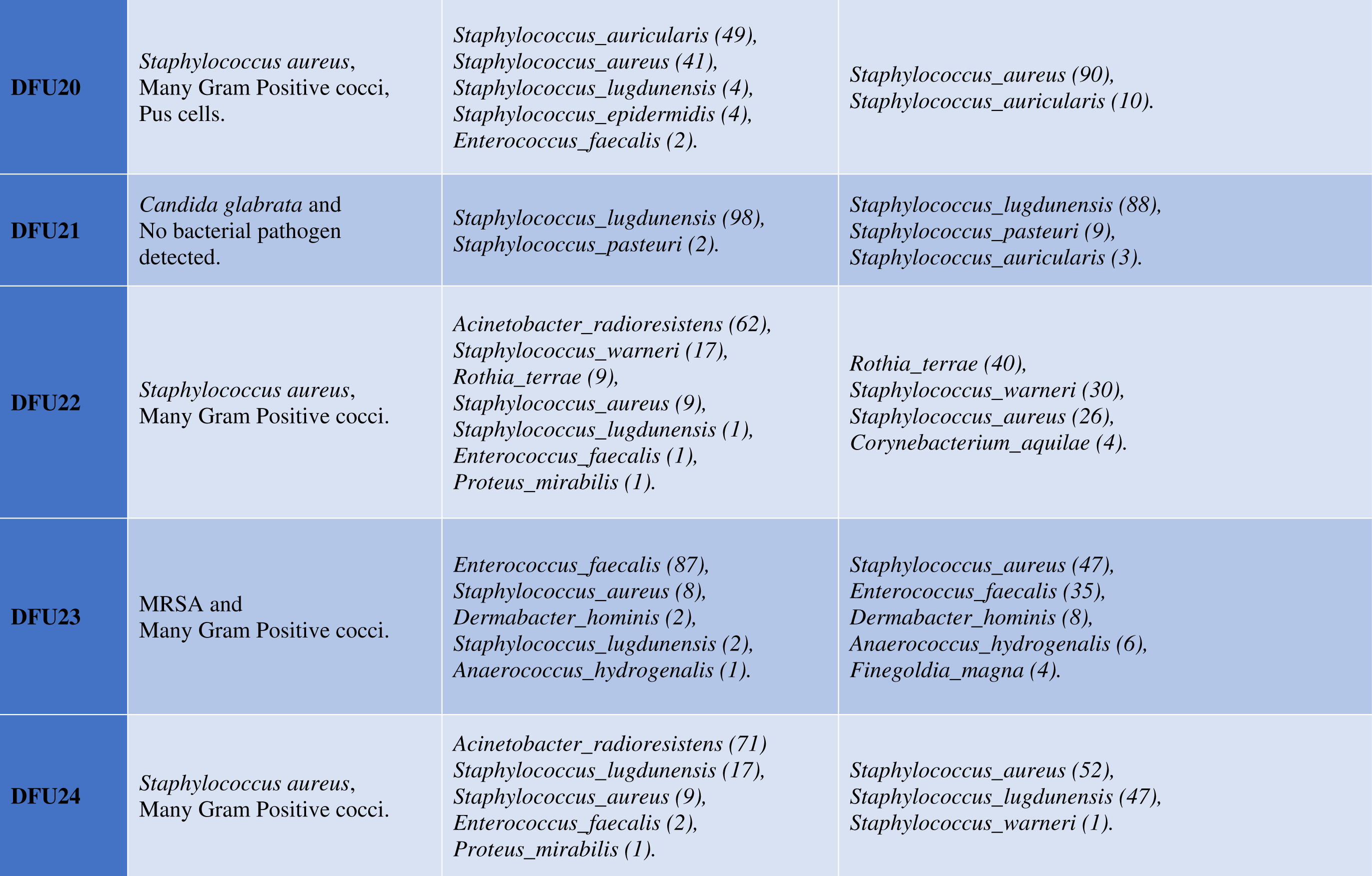
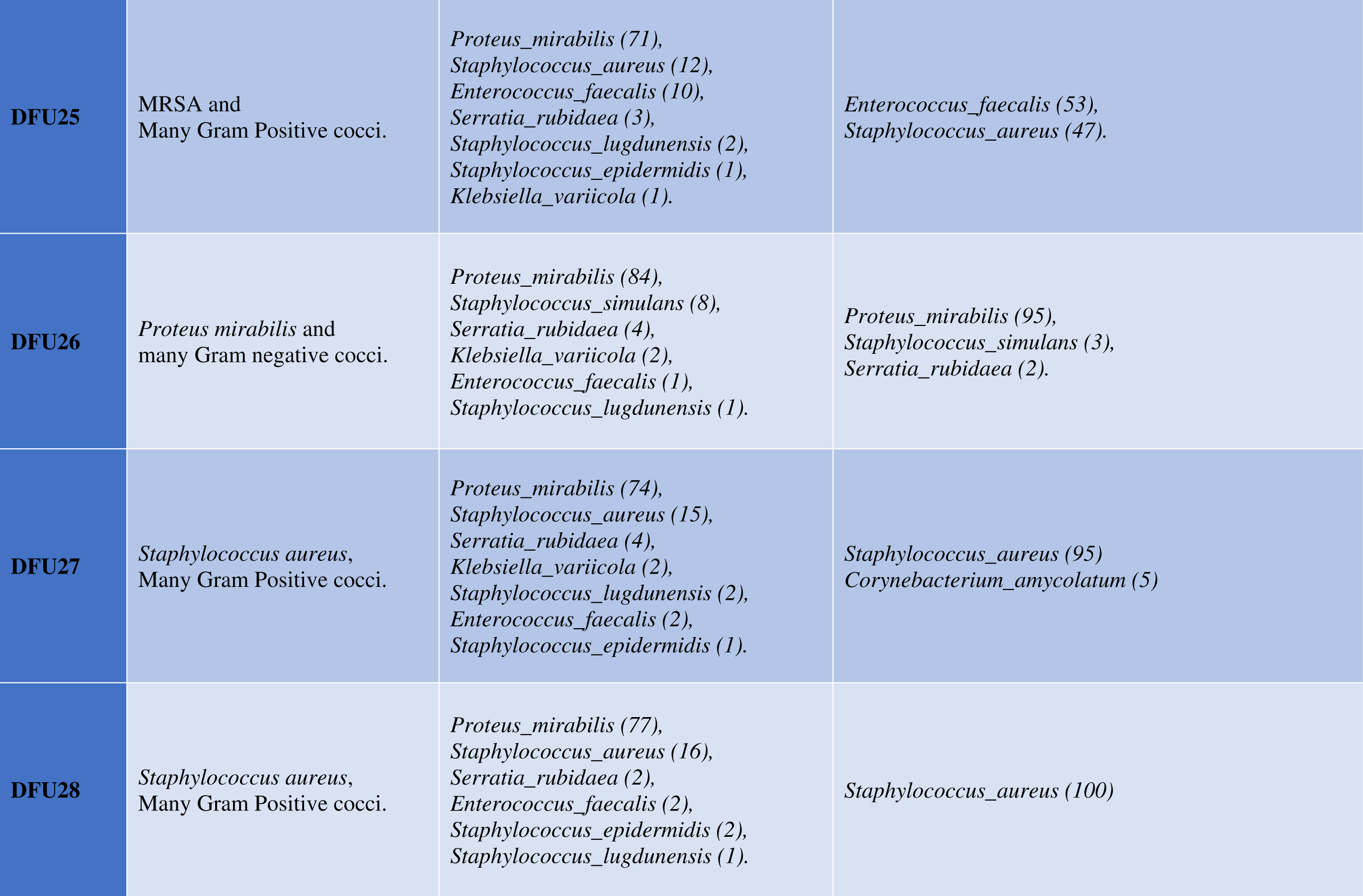
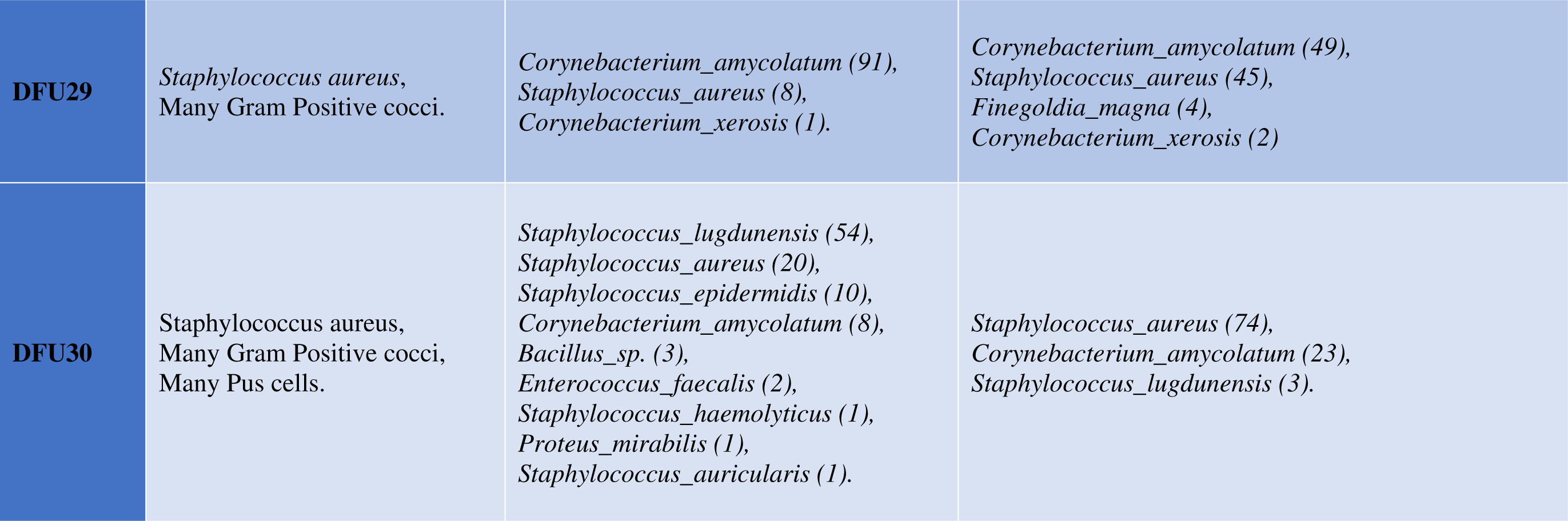
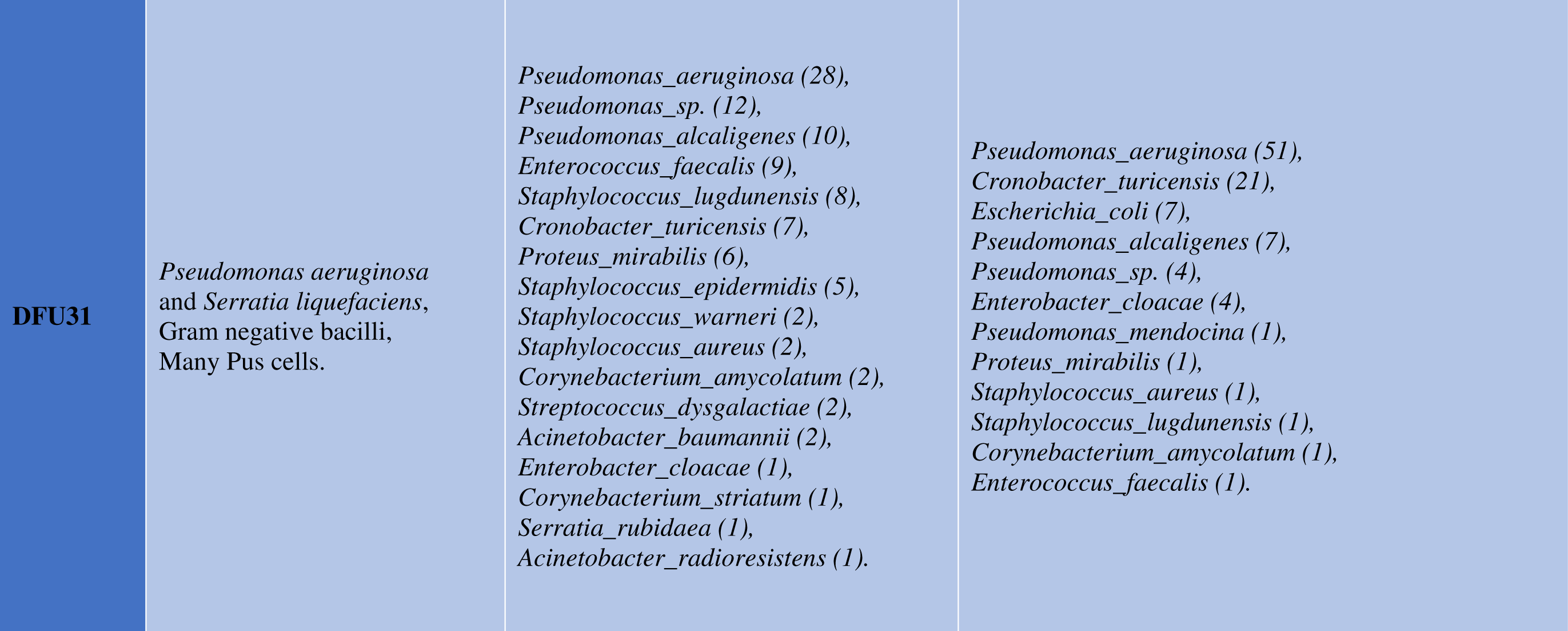
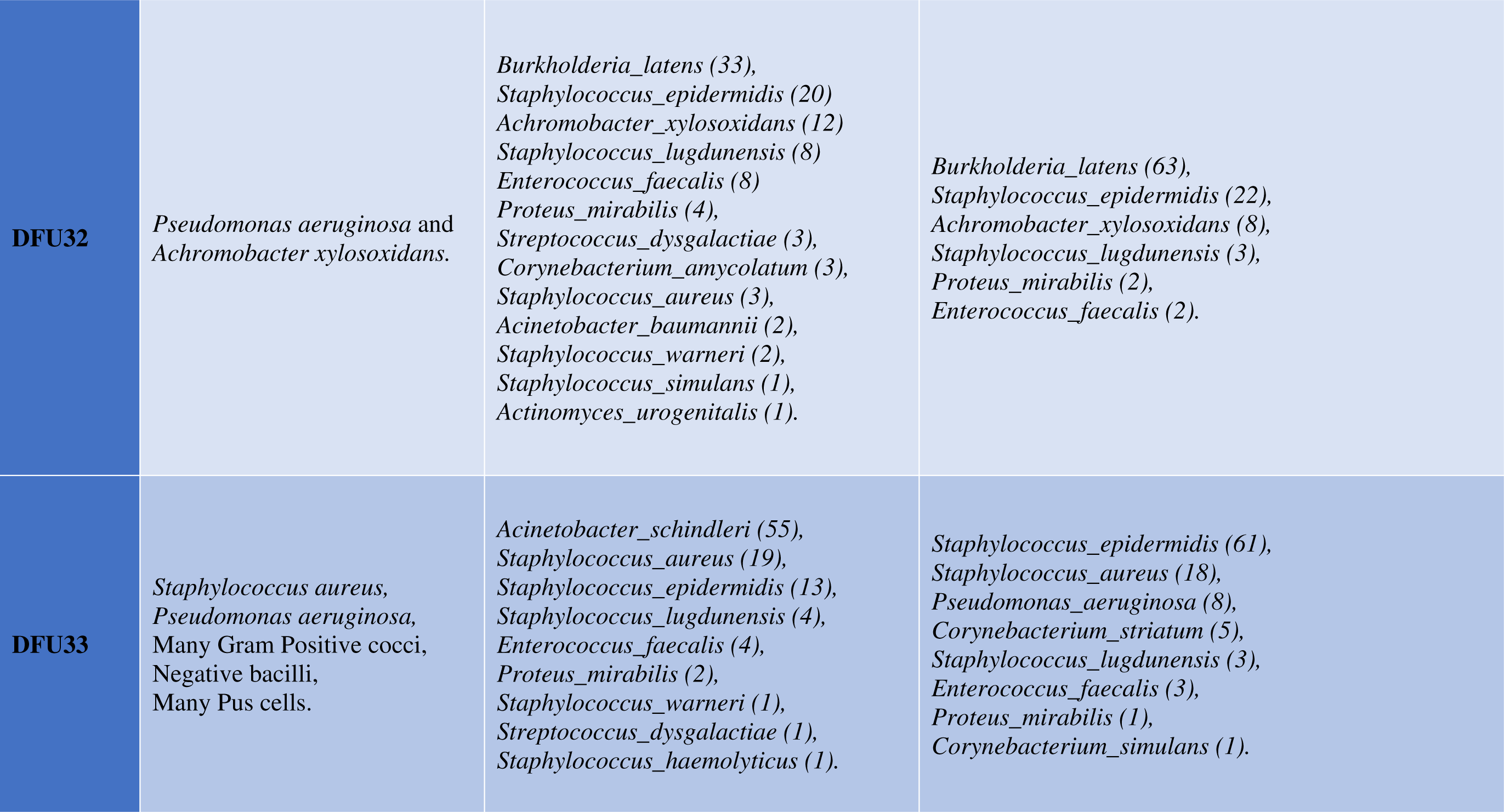
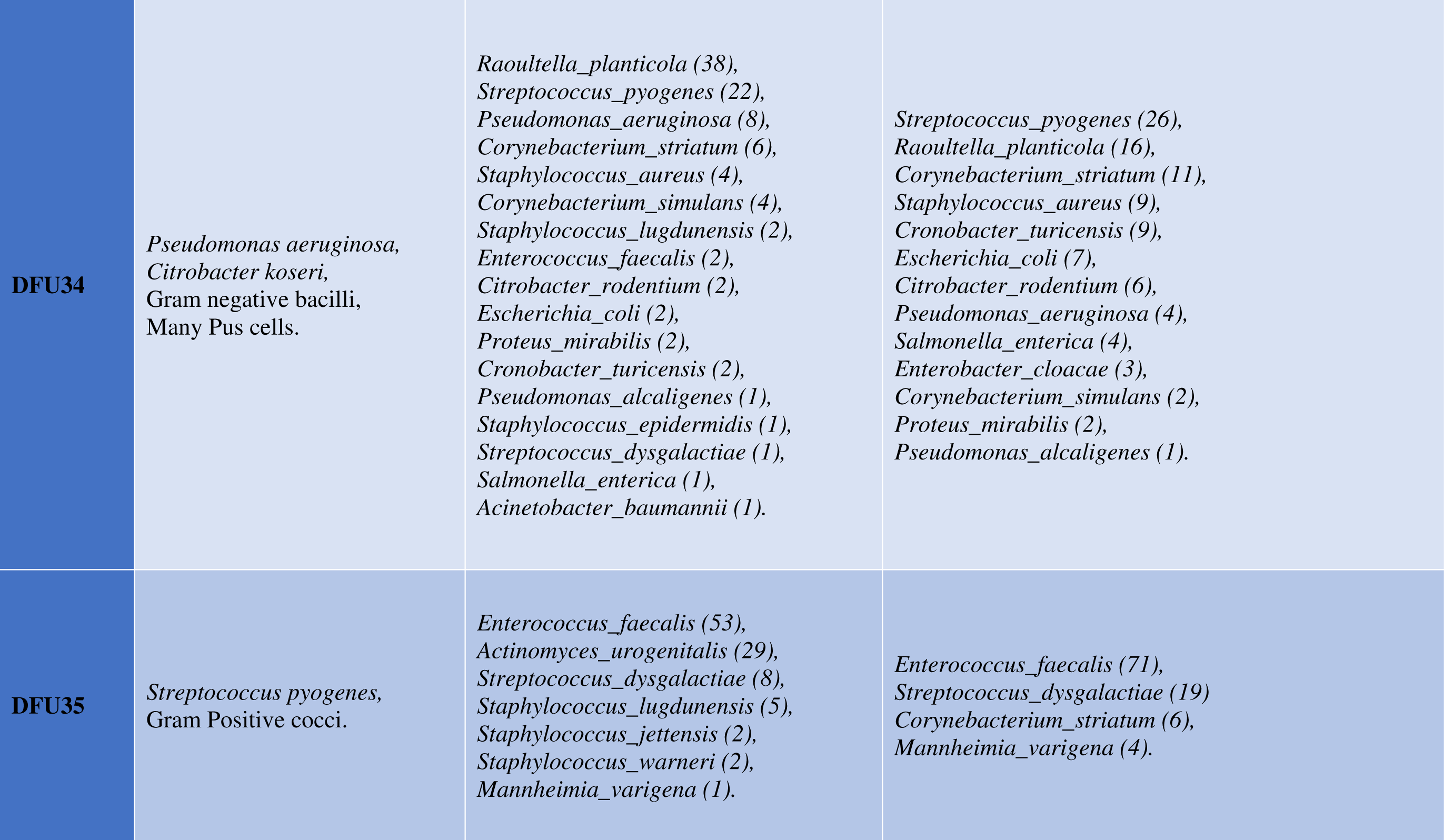
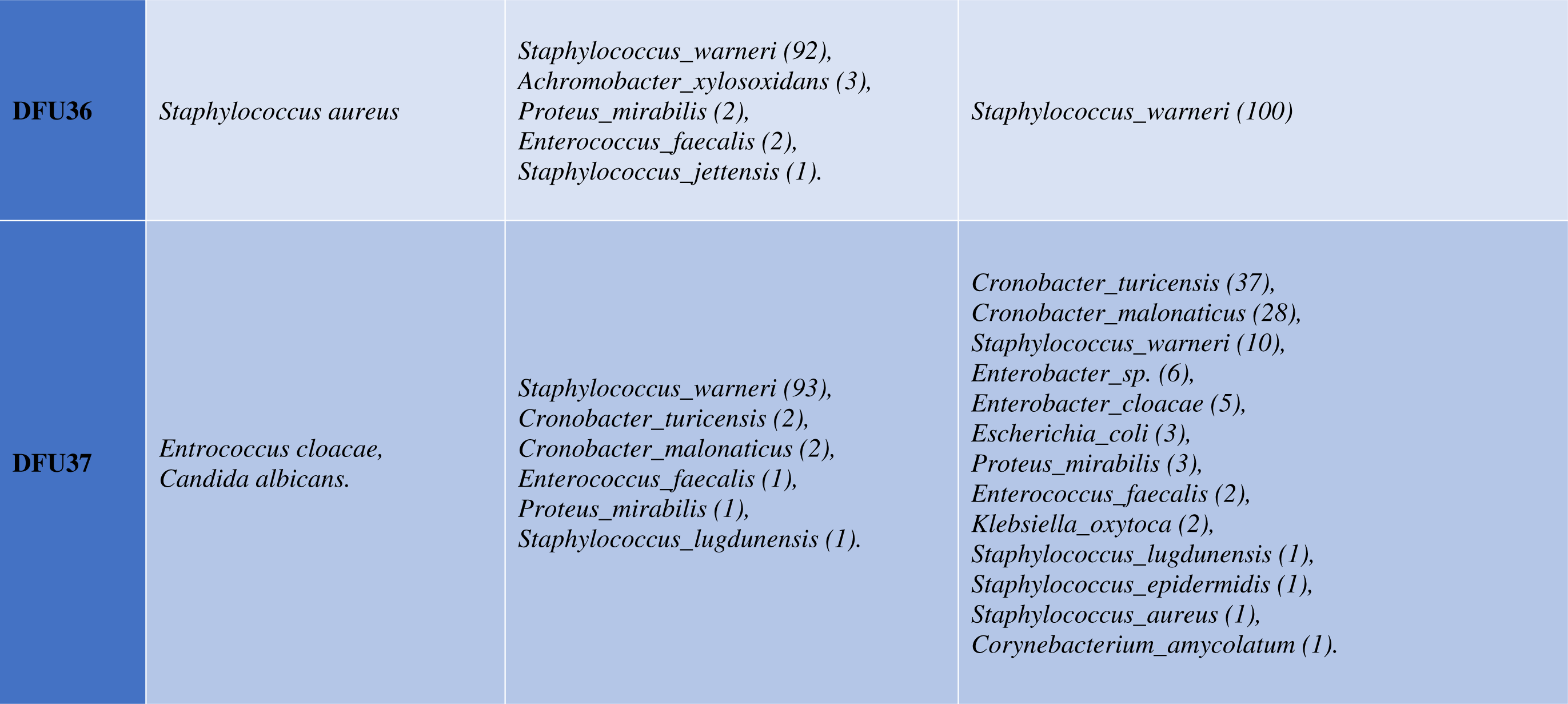
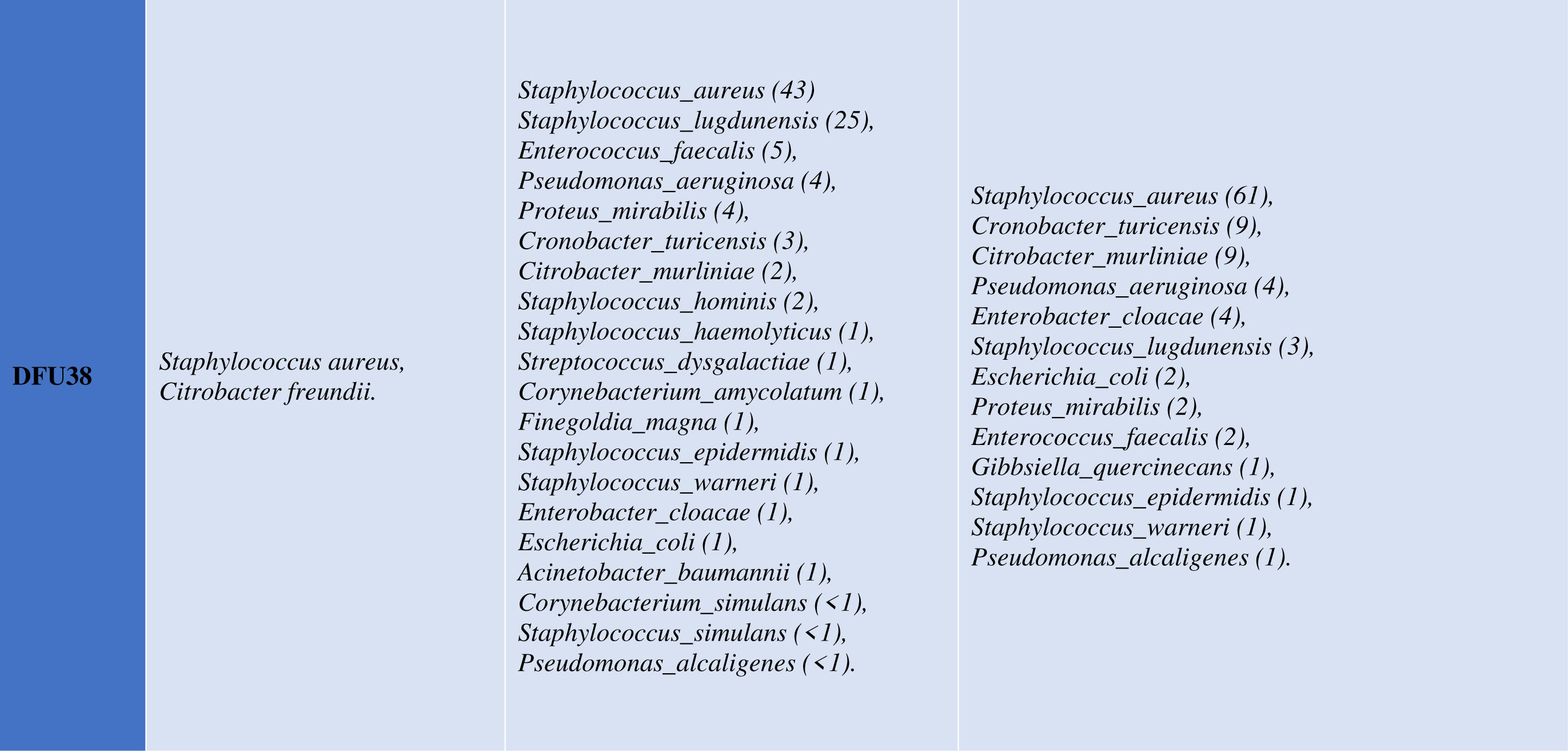
Comparison between different diagnostic methods core DFU microbiome.

Surprisingly, *Pseudomonas aeruginosa,* the well-established DFU pathogen, represented the highest fraction of the core DFU microbiome in only one case (DFU 31) (2.6%) for both microbiome methods. The microbiome methods were able to present *Staphylococcus lugdunensis* as a new DFU causal agent since it represented the highest fraction of the core DFU microbiome in 5 cases (13%) in the CM method and in 2 cases (5%) using the DSM method. *S. lugdunensis* was part of the core microbiome in 32 cases (84%) for the CM method and 24 cases (63%) for the DSM method. Besides, *Staphylococcus epidermidis* should not be overlooked as a potential DFU pathogen since it appeared as the highest fraction of the core DFU microbiome in 5 cases (13%) using the DSM method and in three cases (8%) using the CM method. *S. epidermidis* was part of the core microbiome in 21 cases (55%) for the CM method and 18 cases (47%) for the DSM method.

Also, *Staphylococcus warneri* represented the highest fraction of the core DFU microbiome in 2 cases (5%) in the CM method, while it represented the highest fraction of the core DFU microbiome in one case (2.6%) using the DSM method. However, *S. warneri* was part of the core microbiome in 15 cases (39%) for the CM method and 12 cases (31.6%) for the DSM method. Furthermore, among several detected *Corynebacterium species, Corynebacterium striatum* and *Corynebacterium amycolatum* are important DFU pathogens. *C. striatum* represented the highest fraction of the core DFU microbiome in one case (8%) in the CM method. Moreover, *C. striatum* was part of the core microbiome in 5 cases (13%) for the CM method and in 6 cases (15.8%) for the DSM method. For *C. amycolatum,* it appeared as the highest fraction of the core DFU microbiome in one case (8%) in the CM method and in 2 cases (5%) for the DSM method. It was part of the core microbiome in 10 cases (26%) for the CM method and 14 cases (36.8%) for the DSM method. Moreover, one important pathogen is *Acinetobacter radioresistens* that requires close attention. It appeared as the highest fraction of the core DFU microbiome in three cases (8%) in the CM method. *A. radioresistens* was part of the core microbiome in 4 cases (10%) for the CM method.

### 3.9 Microbial and clinical description of bacterial species detected by CM and DSM methods

Figure 1, together with the S2 table, represent basic Microbial and clinical description of bacterial species detected by CM and DSM methods. Among the 176 OTUs/species, 81 (46%) were Gram-negatives and 95 (54%) Gram positives. 48 OTUs/species were aerobes, five anaerobes, 103 facultative anaerobe, 14 microaerophile, two obligate aerobe, and four obligate anaerobes. Among the 176 OTUs/species, 115 are clinically significant taxa, 33 were clinically insignificant, 3 with questionable clinical significance, and 25 with no information about their clinical significance (**Figure 4**). Fifty-three taxa belong to class one German biosafety level, 93 belong to class two, and 30 with unknown classification.

**Figure 4.**
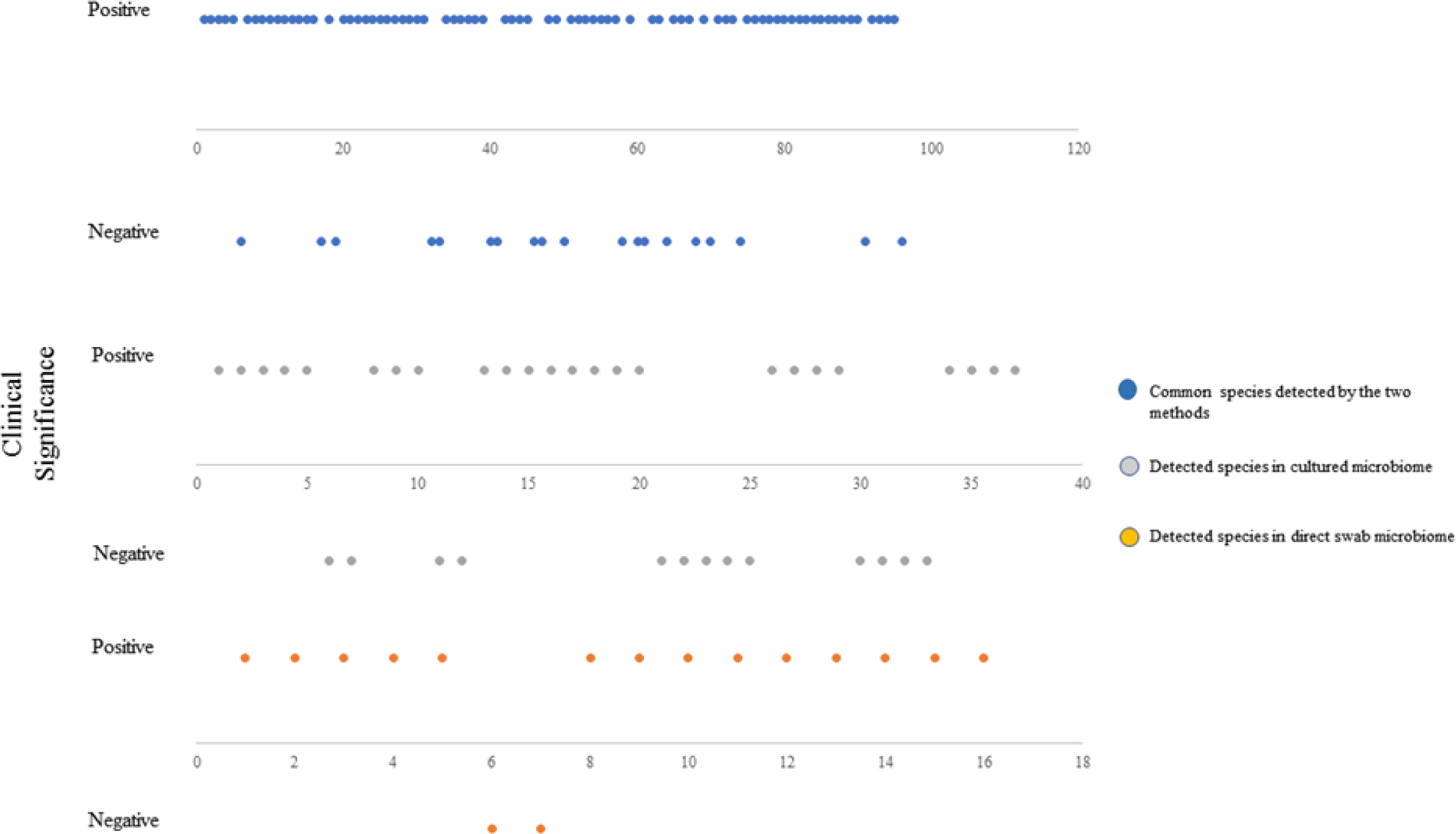
Clinical significance of 176 OTUs/species detected by CM and DSM methods

### 3.10 Skin, wound, or bone pathogenicity of CM and DSM methods specific taxa

Supplementary Table S3 represents the 40 detected species exclusive taxa detected in the CM method only. Out of the 40 taxa, 15 were known skin, wound, or Bone pathogens (37.5%). Supplementary Table S4 represents the 21 detected species exclusive taxa detected in DSM method only. Out of the 21 taxa, ten were known skin, wound, or Bone pathogens (47.6%).

### 3.10 Bacterial Relative abundance distribution amongst Neuropathic and Neuroischaimic DFU using the CM method

#### 3.10.1 Relative abundance distribution of *Staphylococcus* species

As shown in the S1 table, our 38 study subjects are divided into 26 subjects with neuroischaimic DFU (ISCDFU) and 12 subjects with neuropathic DFU (NURDFU). **Figure 5** presents the relative abundance of the different observed *Staphylococcus* species in both types of ulcers. Generally, we observed the presence of 24 species/OTUs belonging to genus Staphylococcus, namely, *S. agnetis, S. aureus, S. auricularis, S. capitis, S. chromogenes, S. cohnii, S. devriesei, S. epidermidis, S. equorum, S. haemolyticus, S. hominis, S. jettensis, S. lugdunensis, S. pasteuri, S. petrasii, S. pettenkoferi, S. piscifermentans, S. pseudolugdunensis, S. saccharolyticus, S. saprophyticus, S. sciuri, S. simulans, Staphylococcus sp., and S. warneri.* However, *S. devriesei, S. equorum*, *S. pseudolugdunensis,* and *S. sciuri* were not present in ISCDFU. On the other hand, *S. capitis* and *S. saprophyticus* were not present in NURDFU. The highest total relative abundance in ISCDFU was observed with *S. warneri* (30%), followed by *S. aureus* and *S. lugdunensis* (26%), then *S. epidermidis* (10%). On the other hand, the highest total relative abundance in NURDFU was observed with *S. lugdunensis* (58%), followed by *S. aureus* (16%), then *S. epidermidis* (9%).

**Figure 5.**
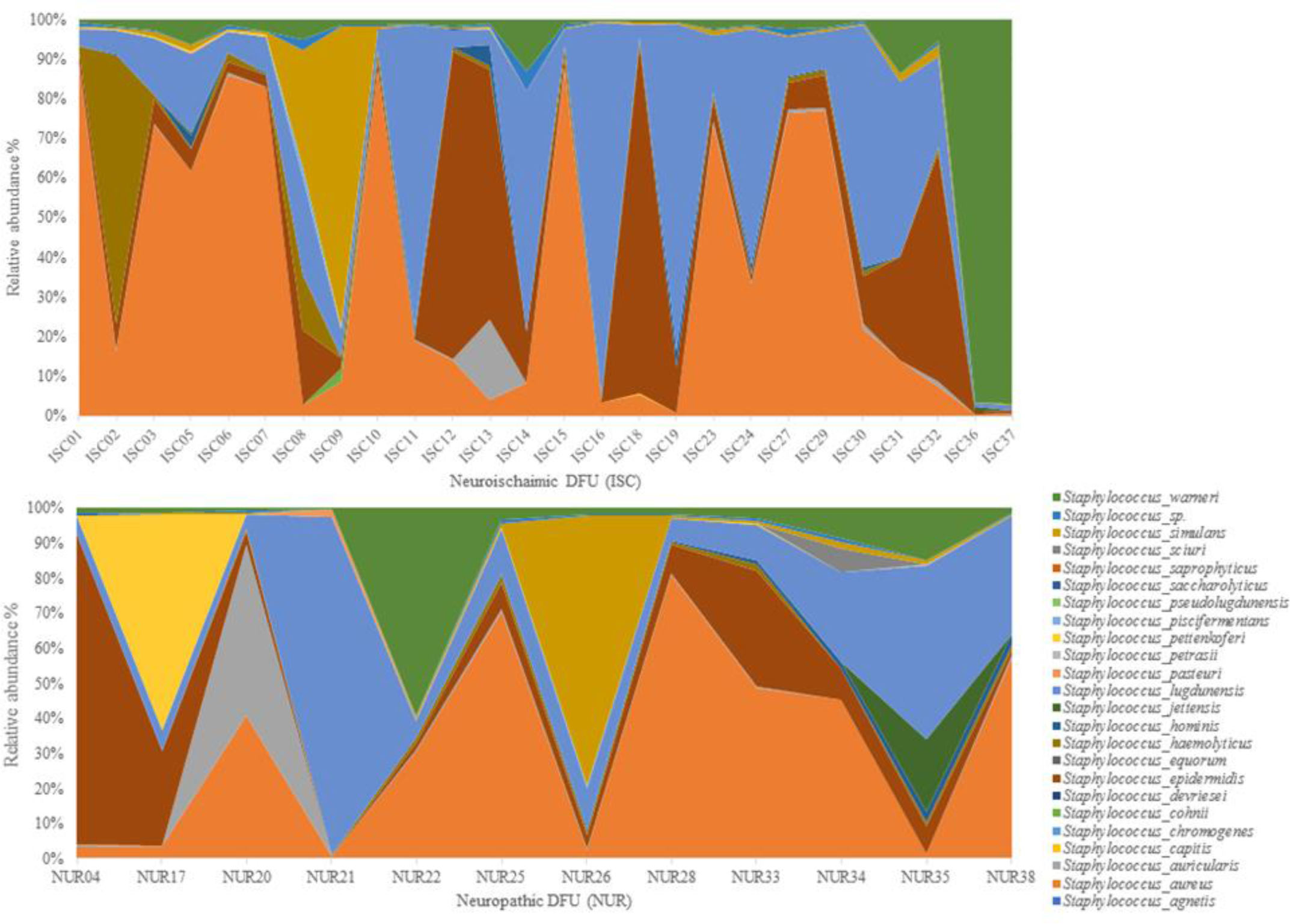
Relative abundance distribution of *Staphylococcus* species among individuals with ISCDFU and NURDFU

#### 3.10.2 Relative abundance distribution of *Acinetobacter* species

**Figure 6** presents the relative abundance of the different observed *Acinetobacter* species in both types of ulcers. Generally, we observed the presence of ten species/OTUs belonging to the genus Acinetobacter, namely, *A. baumannii, A. guillotine, A. johnsonii, A. junii, A. lwoffii, A. radioresistens, A. rhizosphaerae, A. schindleri, A. soli,* and *Acinetobacter sp.* However, *A. guillouiae, A. johnsonii*, and *A. lwoffii* were not present in ISCDFU. On the other hand, *A. rhizosphaerae* was not present in NURDFU. The highest total relative abundance in ISCDFU was observed with *A. baumannii* (56%), followed by *A. radioresistens* (42%). On the other hand, the highest total relative abundance in NURDFU was observed with *A. radioresistens* (78%), followed by *A. baumannii* (18%).

**Figure 6.**
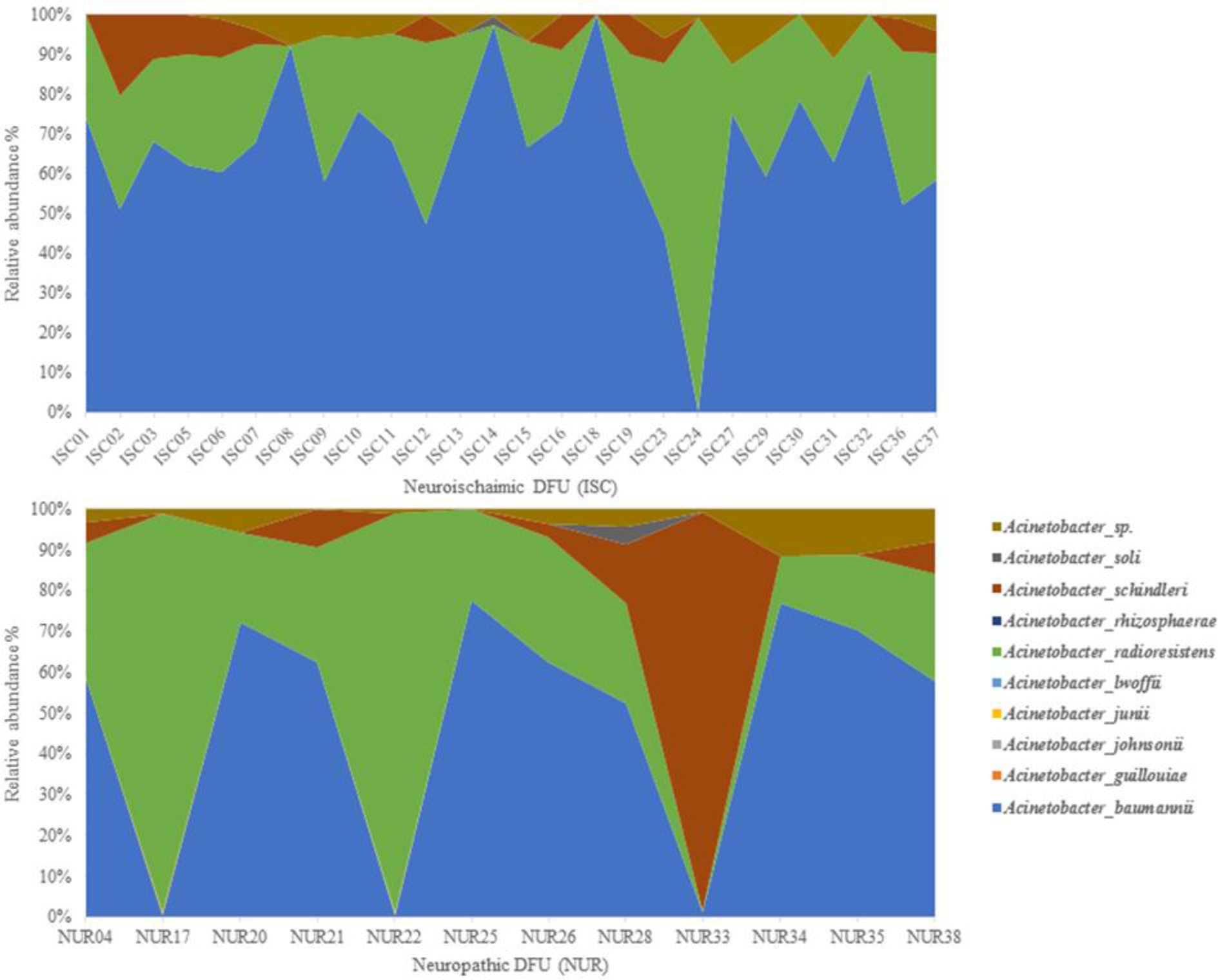
Relative abundance distribution of *Acinetobacter* species among individuals with ISCDFU and NURDFU

#### 3.10.3 Relative abundance distribution of *Corynebacterium* species

**Figure 7** presents the relative abundance of the different observed *Corynebacterium* species in both types of ulcers. Generally, we observed the presence of 12 species/OTUs belonging to the genus Corynebacterium, namely, *C. afermentans, C. amycolatum, C. aquilae, C. aurimucosum, C. auriscanis, C. jeikeium, C. mucifaciens, C. simulans, C. striatum, C. ureicelerivorans, C. xerosis,* and *Corynebacterium sp.* However, *C. aquilae* and *C. aurimucosum* were not present in ISCDFU. On the other hand, *C. afermentans, C. auriscanis, C. jeikeium, C. mucifaciens,* and *C. ureicelerivorans* were not present in NURDFU. The highest total relative abundance in ISCDFU was observed with *C. amycolatum* (68%), followed by *C. striatum* (17%), then *C. simulans* (13%). On the other hand, the highest total relative abundance in NURDFU was observed with *C. amycolatum* (40%), followed by *C. striatum* (35%), then *C. simulans* (18%).

**Figure 7.**
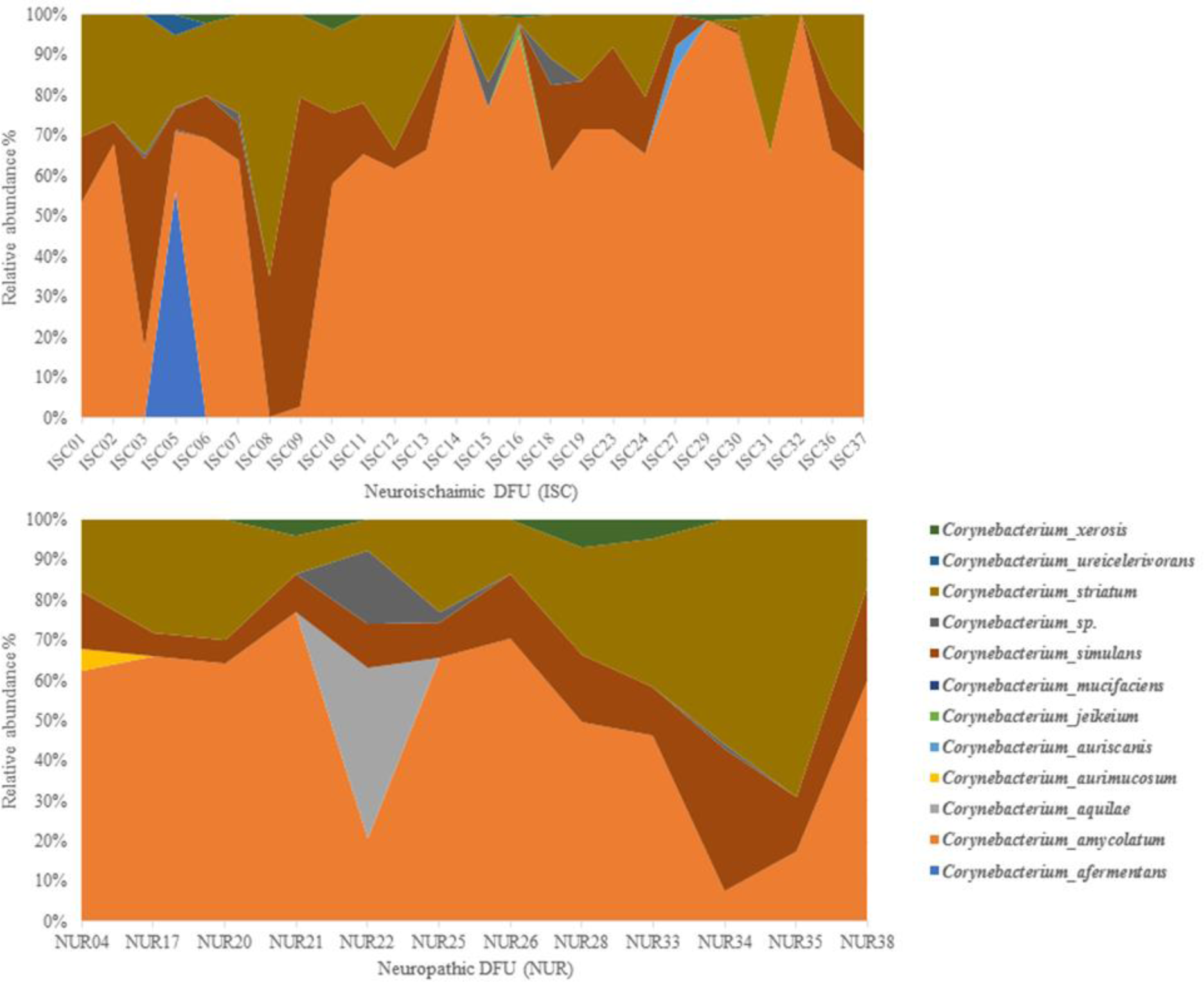
Relative abundance distribution of *Corynebacterium* species among individuals with ISCDFU and NURDFU

#### 3.10.4 Relative abundance distribution of *Pseudomonas* species

**Figure 8** presents the relative abundance of the different observed *Pseudomonas* species in both types of ulcers. Generally, we observed the presence of eight species/OTUs belonging to the genus *Pseudomonas*, namely, *P. aeruginosa, P. alcaligenes, P. indica, P. mendocina, P. nitroreducens, P. pohangensis, P. stutzeri,* and *Pseudomonas sp.* However, *P. pohangensis* was not present in ISCDFU. However, *P. indica, P. mendocina,* and *P. nitroreducens* were not observed in NURDFU. The highest value of relative abundance in ISCDFU was observed with *P. aeruginosa* (52%), followed by *P. alcaligenes* (25%), then *Pseudomonas sp.* (21%). On the other hand, the highest total relative abundance in NURDFU was observed with *P. aeruginosa* (79%), followed by *P. alcaligenes* (10%), then *Pseudomonas sp.* (9%).

**Figure 8.**
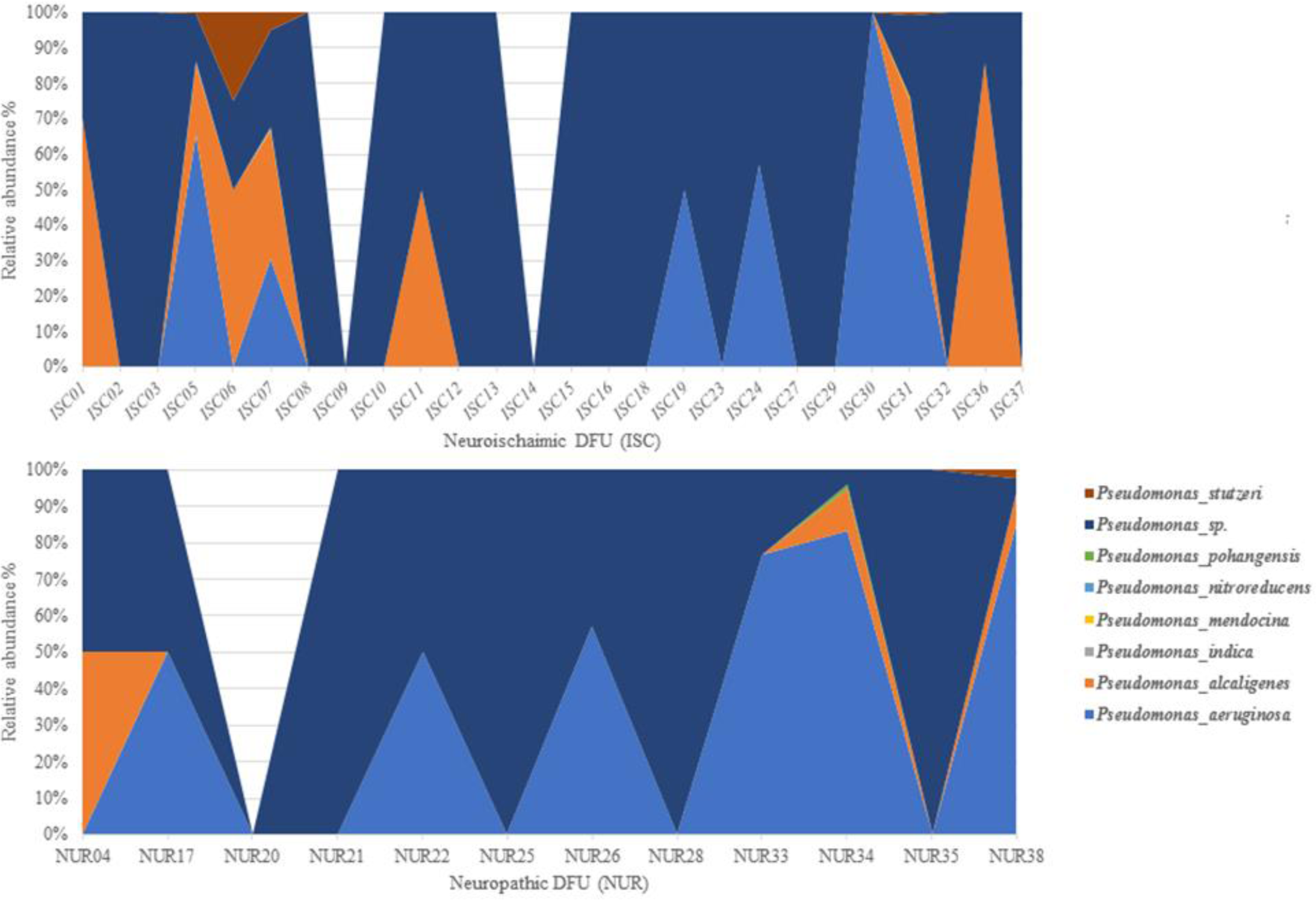
Relative abundance distribution of *Pseudomonas* species among individuals with ISCDFU and NURDFU

#### 3.10.5 Relative abundance distribution of *Streptococcus* species

**Figure 9** presents the relative abundance of the different observed *Streptococcus* species in both types of ulcers. Generally, we observed the presence of 12 species/OTUs belonging to the genus Streptococcus, namely, *S. agalactiae, S. australis, S. dysgalactiae, S. infantis, S. intermedius, S. lactarius S. mitis, S. phocae, S. pneumoniae, S. pyogenes, S. salivarius,* and *Streptococcus sp.* All ten *Streptococcus* species were observed in the ISCDFU. *S. australis, S. infantis, S. intermedius, S. lactarius, S. mitis, S. pneumoniae,* and *S. salivarius* were not observed in the NURDFU. The highest total relative abundance in ISCDFU was observed with *S. dysgalactiae* (99%). On the other hand, the highest total relative abundance in NURDFU was observed with *S. dysgalactiae* (68%), followed by *S. pyogenes* (27%) than *S. phocae* (4%).

**Figure 9.**
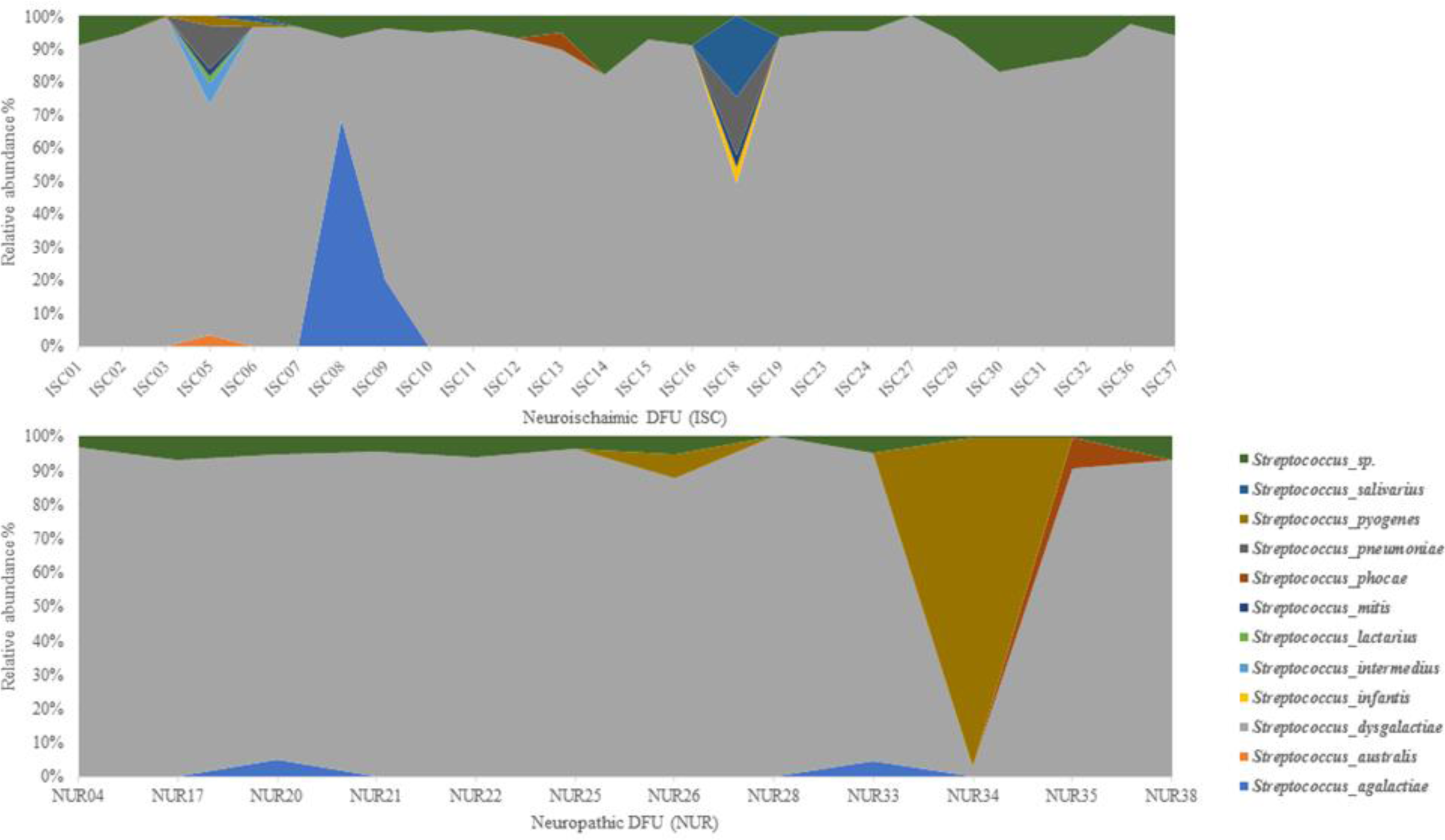
Relative abundance distribution of *Streptococcus* species among individuals with ISCDFU and NURDFU

#### 3.10.6 Relative abundance distribution of *Achromobacter* species

**Figure 10** presents the distribution of the relative abundance of the different observed *Achromobacter* species in both types of ulcers. Generally, we observed the presence of three species/OTUs belonging to the genus Achromobacter, namely, *A. insolitus, A. xylosoxidans,* and *Achromobacter sp.* The highest total relative abundance in ISCDFU was observed with *A. xylosoxidans* (89%) followed by *Achromobacter sp.* (9%), then *A. insolitus* (2%). On the other hand, the highest total relative abundance in NURDFU was observed with *Achromobacter sp.* (64%), followed by *A. xylosoxidans* (25%), then *A. insolitus* (11%).

**Figure 10.**
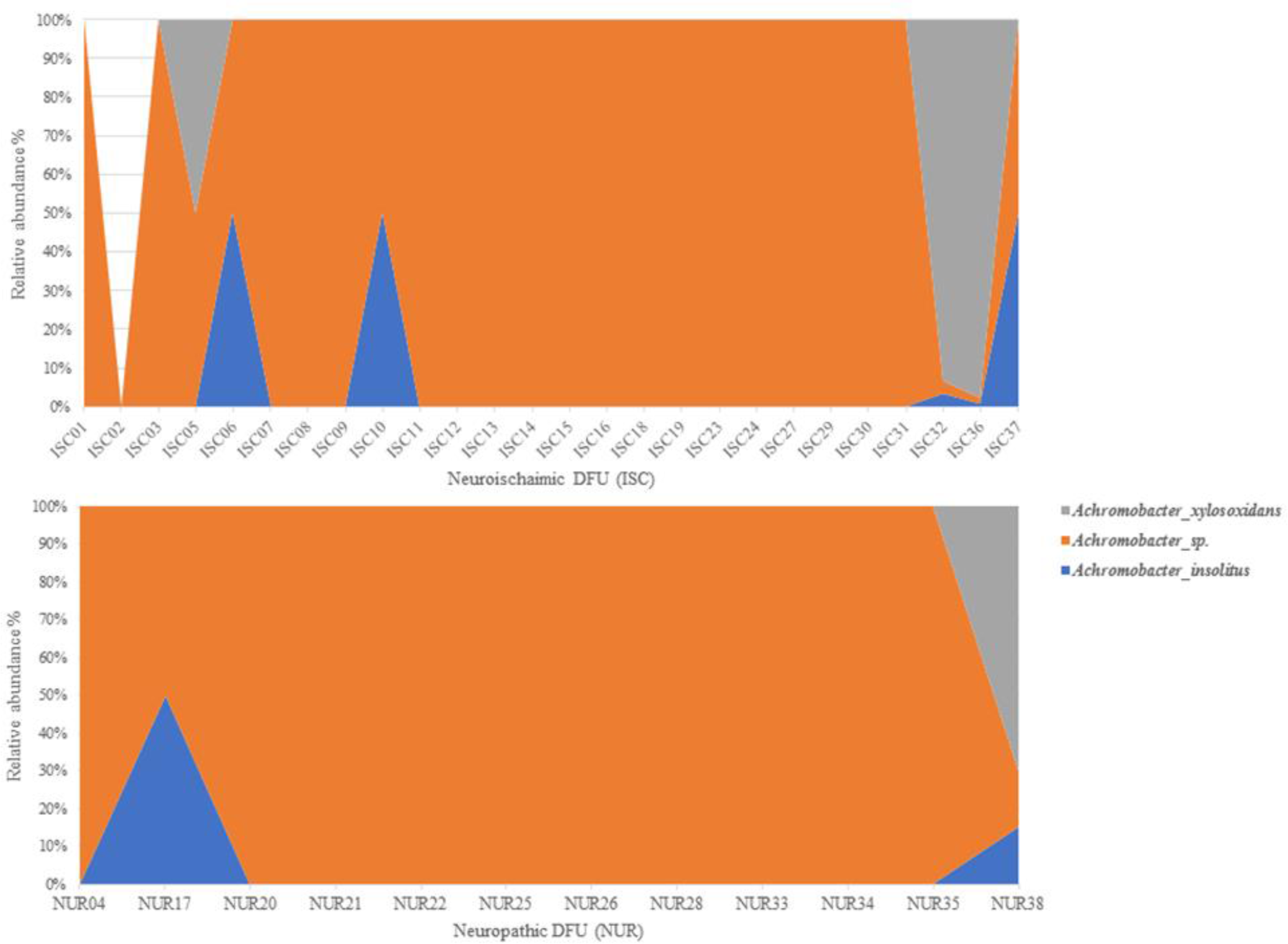
Relative abundance distribution of *Achromobacter* species among individuals with ISCDFU and NURDFU

#### 3.10.7 Relative abundance distribution of other observed bacterial species

We observed the presence of five species/OTUs belonging to the genus *Serratia* namely, *S. marcescens, S. plymuthica, S. rubidaea, S. symbiotica,* and *Serratia sp*. The highest total relative abundance in ISCDFU was observed with *S. rubidaea* (79%), followed by *S. marcescens* (18%). The highest total relative abundance in NURDFU was observed with *S. rubidaea* (84 %), followed by *S. marcescens* (13%). Moreover, we observed the presence of two species/OTUs belonging to the genus *Proteus* namely, *P. mirabilis* and unresolved *Proteus sp. P. mirabilis* (>99.9%) represented the highest total relative abundance in both types of ulcer. Besides, we observed the presence of 8 species/OTUs belonging to genus *Enterococcus* namely, *E. avium, E. casseliflavus, E. faecalis, E. faecium, E. gallinarum, E. lactis, E. lemanii,* and *Enterococcus sp. E. faecalis* represented the highest total relative abundance in ISCDFU and NURDFU, 99.9% and 96% respectively. Moreover, we observed the presence of three species/OTUs belonging to the genus *Actinomyces*, namely, *A. neuii, Actinomyces sp.,* and *A. urogenitalis*. *A. urogenitalis* represented the highest total relative abundance in ISCDFU and NURDFU, 90% and 99.9%, respectively. Additionally, we observed the presence of four species/OTUs belonging to the genus *Klebsiella* namely, *K. oxytoca, K. pneumoniae, Klebsiella sp.,* and *K. variicola. K. variicola* represented the highest total relative abundance in ISCDFU and NURDFU, 64% and 74%, respectively. Furthermore, we observed the presence of two species/OTUs belonging to genus *Brevibacterium*, namely, *B. paucivorans* and unresolved *Brevibacterium sp*. *B. paucivorans* represented the highest total relative abundance in ISCDFU and NURDFU, 99.9% and 100% respectively. Moreover, we observed the presence of seven species/OTUs belonging to the genus *Bacillus,* namely, *B. cereus, B. licheniformis, B. pumilus, B. seohaeanensis, B. thermophilus, B. thuringiensis, and Bacillus sp.* Unresolved *Bacillus sp.* showed the highest total relative abundance in ISCDFU at 87%, while *Bacillus sp.* and *B. seohaeanensis* showed the highest total relative abundance values NURDFU 41% and 38%, respectively. Furthermore, we observed two species/OTUs belonging to genus *Dermabacter,* namely, *D. hominis* and *Dermabacter sp. D. hominis* showed the highest total relative abundance in ISCDFU and NURDFU, 99.9% and 100%, respectively. Furthermore, we observed *Finegoldia magna, Staphylococcus saccharolyticus,* and *Morganella morganii* in both types of ulcers. Moreover, we observed *Anaerococcus hydrogenalis* only in ISCDFU and *Bacteroides fragilis* and *Bacteroides sp.* only in NURDFU.

#### 3.11 Biological diversity comparison between Neuropathic and Neuroischaimic DFU

**S5 and S6 tables** present the diversity indices for the 26 subjects with neuroischaimic DFU (ISCDFU) and 12 subjects with neuropathic DFU (NURDFU), respectively. Generally, the NURDFU group showed higher values for the tested diversity indices than the ISCDFU group. The number of observed taxa ranged between 19-66 with a mean of 34.5769 taxa with a sum of 899 for the ISCDFU group. However, the number of observed taxa ranged between 32-51 with a mean of 41 taxa with a sum of 492 for the NURDFU group. The Simpson 1-D index ranged between 0.1291-0.8813 with a mean of 0.4568 for the ISCDFU group. Whereas the Simpson 1-D index ranged between 0.1008- 0.8114 with a mean of 0.5669 for the NURDFU group. The Shannon-H index ranged between 0.3795- 2.568 with a mean of 1.1887 for the ISCDFU group. However, the Shannon-H index ranged between 0.3208-2.28 with a mean of 1.4021 for the NURDFU group. The Margalef index ranged between 1.92-6.793 with a mean value of 3.696 for the ISCDFU group. While the Margalef index ranged between 3.098- 5.567 with a mean value of 4.3493 for the NURDFU group. Finally, the Chao-1 index ranged between 19-66 with a mean of 34.5769 for the ISCDFU group. Whereas the Chao-1 index ranged between 32-51 with a mean of 41 taxa. Only Taxa-s, Chao-1, and Margalef indices were significantly different between the two types of Ulcers (**Figures 11**).

**Figure 11.**
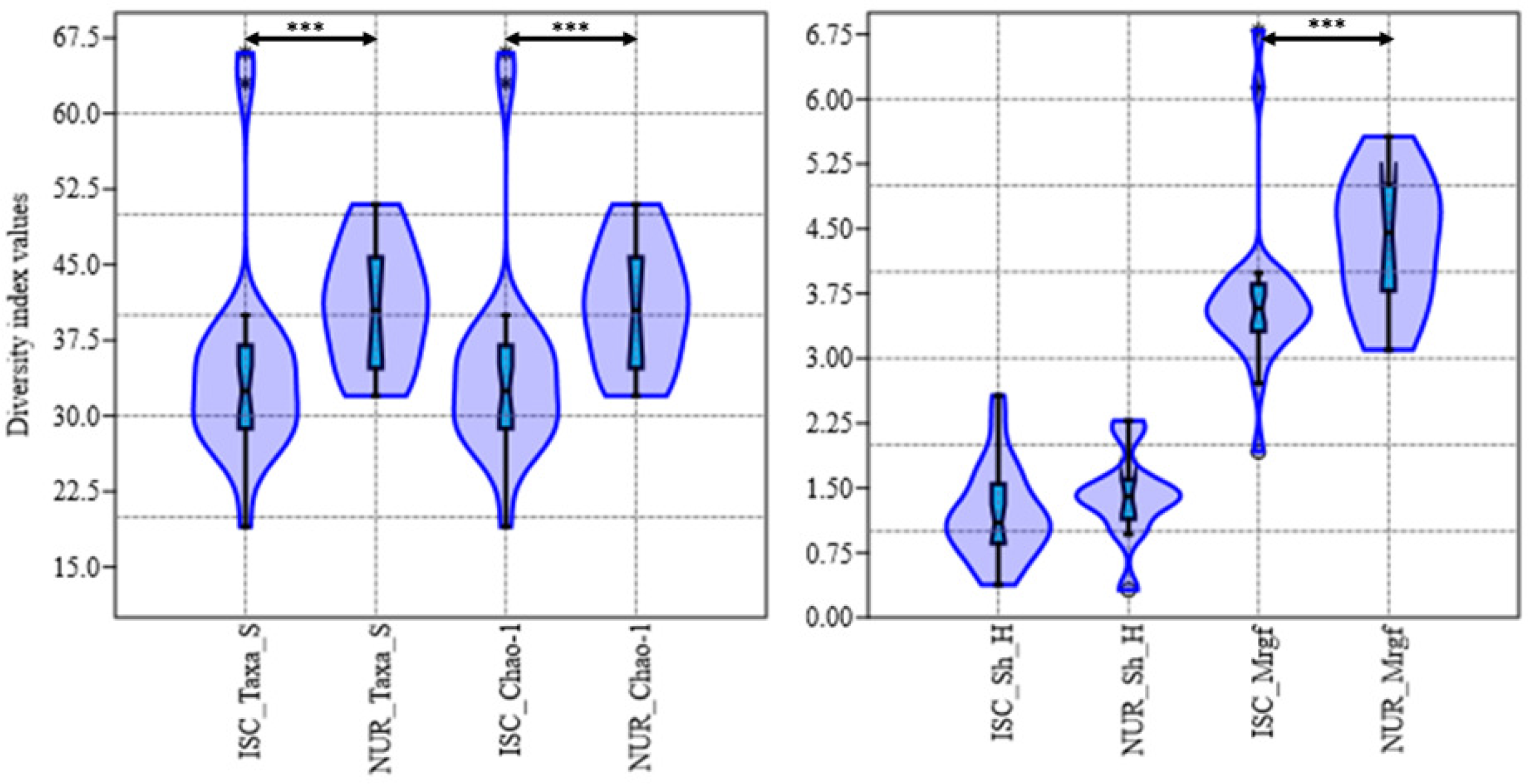
Violin plots of the diversity indices Taxa-s, Chao-1, Shannon H (Sh-H), and Margalef (Mrgf) for two types of Ulcers.

#### 3.12 Cluster analysis and Principal component analysis (PCA) for ISCDFU and NURDFU microbiomes

Neither Cluster analysis UPGMA (Correlation, elucidation or Bary-Curtis criteria) (**Figure 12**) or Principal component analysis (PCA) **(Figure 13a and b)**, based on both qualitative and quantitative bacterial distribution, showed no apparent clustering or pattern amongst the two types of ulcers, namely, NURDFU and ISCDFU. The cluster analysis showed the presence of 13 clades/clusters, out of which 6 included the two types of ulcers and the remaining seven presented clades that contain single or multiple ISCDFU microbiomes. Cluster/clade VII contained the largest number of microbiomes (8) divided into 5 ISCDFU and 3 NURDFU microbiomes. On the other hand, Clades II, IIII, VIIII, XII, and XIII included only one distinct ISCDFU microbiome. The NURDFU microbiomes always shared clades with ISCDFU microbiomes with clade VI containing three NURDFU microbiomes. Generally, the investigated clinical factors distributed arbitrarily among the observed 13 clusters. For example, the most extensive and mixed ulcer types clade VII, contained values of HbA1C that ranged between 7.8-11.8, diabetes duration between 12-36 years, BMI between 28.78-36.80, wound surface area between (WSA) 1-56 cm^2^, Ankle-brachial pressure index (ABPI) between 0.45-1.29. It also contained individuals/DFU microbiomes associated with only two diabetes complications (Neuropathy and obesity) and individuals/DFU microbiomes with six diabetes complications (Neuropathy, obesity, retinopathy, hypertension, nephropathy, and vasculopathy). Besides, it also contained surface wounds and deep to bone wounds. Clade VII also contained DFU microbiomes belonging to different foot locations, such as the big toe, heel, and planter. Moreover, several ulcer causes were observed in the same clade (VII), such as trauma, footwear, and callus. Furthermore, the ISCDFU clade VIII contained three ISCDFU microbiomes and with no observed NURDFU microbiomes. Similarly, it contained a wide range of some clinical factors such as diabetes duration (7-30 years), HbA1C (9.1-13.4), BMI (21.36-39.24). VIII contained both surface and deep to the tendon wounds, microbiomes associated with only two diabetes complications (Neuropathy and retinopathy), and microbiomes associated with five diabetes complications (Neuropathy, retinopathy, hypertension, obesity, and hyperlipidemia). Clade VIII contained microbiomes of DFUs caused by either callus or footwear and located at the big toe, hallux of the big toe, or the foot’s planter. However, a slimmer range of WSA (1-2 cm^2^) and ABPI (0.32-0.84) was observed. Besides, clade I also contained only two ISCDFU microbiomes, namely, ISC 36 and ISC 37. Similarly, it contained a wide range of clinical factors such as diabetes duration (1-10 years), HbA1C (8.6-13.4). Clade I contained microbiomes of DFUs caused by either trauma or heat and located at the big toe or planter of the foot and either surface or deep to the tendon wounds. However, slimmer range of BMI (31.36-33.75), diabetes complications (4-5), WSA (12 cm^2^), and ABPI (0.49-0.51). Both clade I and clade VIII were supported by relatively higher N1000 bootstrap values of 76 and 59, respectively. Likewise, Principal component analysis (PCA) supported by the minimum spanning tree analysis, which investigates the spatial distribution of a point pattern with a focus on small scales, comparable to nearest neighbor analysis but with somewhat different properties, showed that ISCDFU and NURDFU microbiomes distributed indiscriminately and without any evident clustering both before and after eliminating the after the outliers **(Figure 13a and b)**.

**Figure 12.**
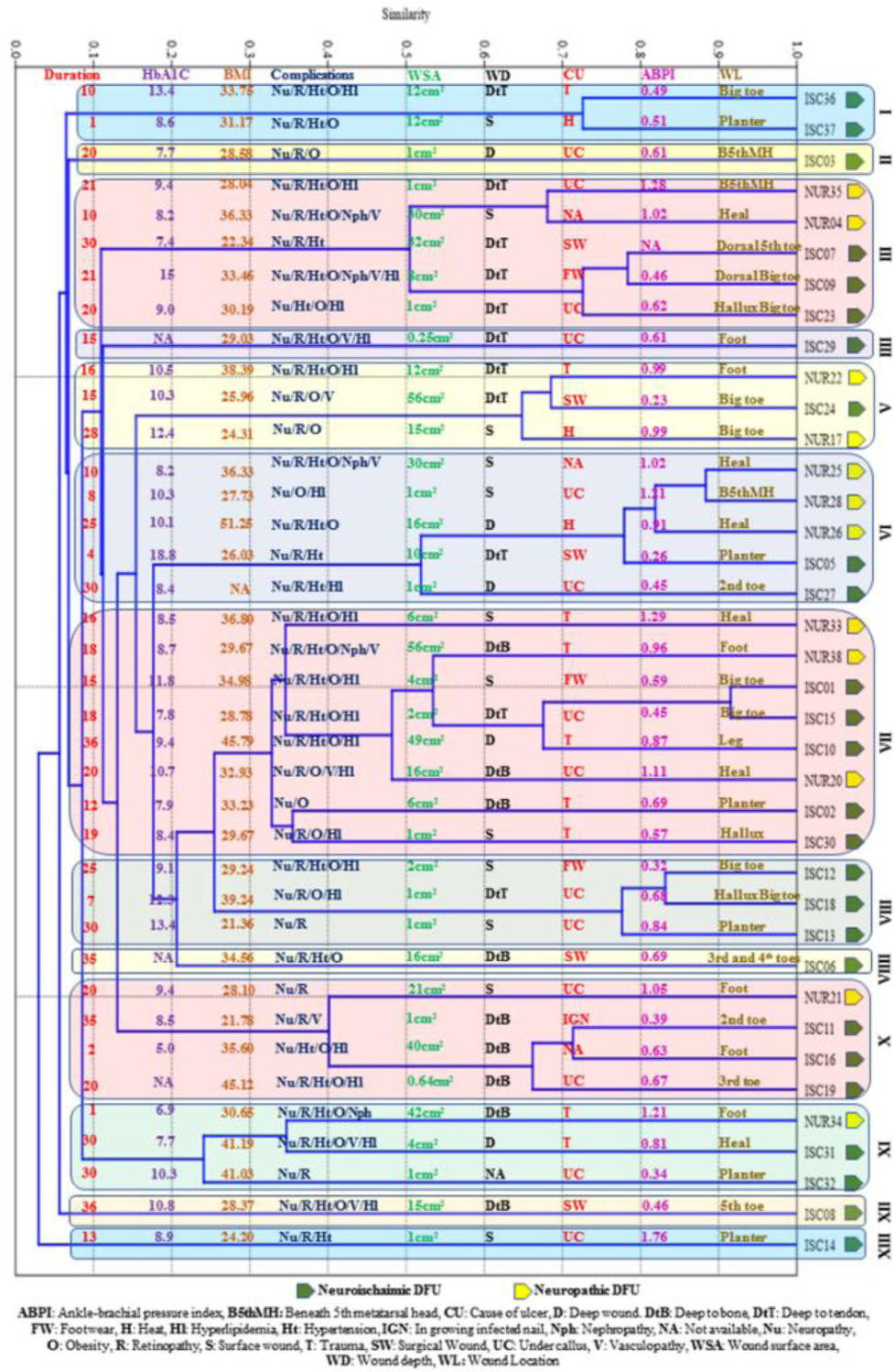
Cluster analysis UPGMA (Correlation criteria) based on both qualitative and quantitative bacterial distribution in the DFU microbiome

**Figure 13.**
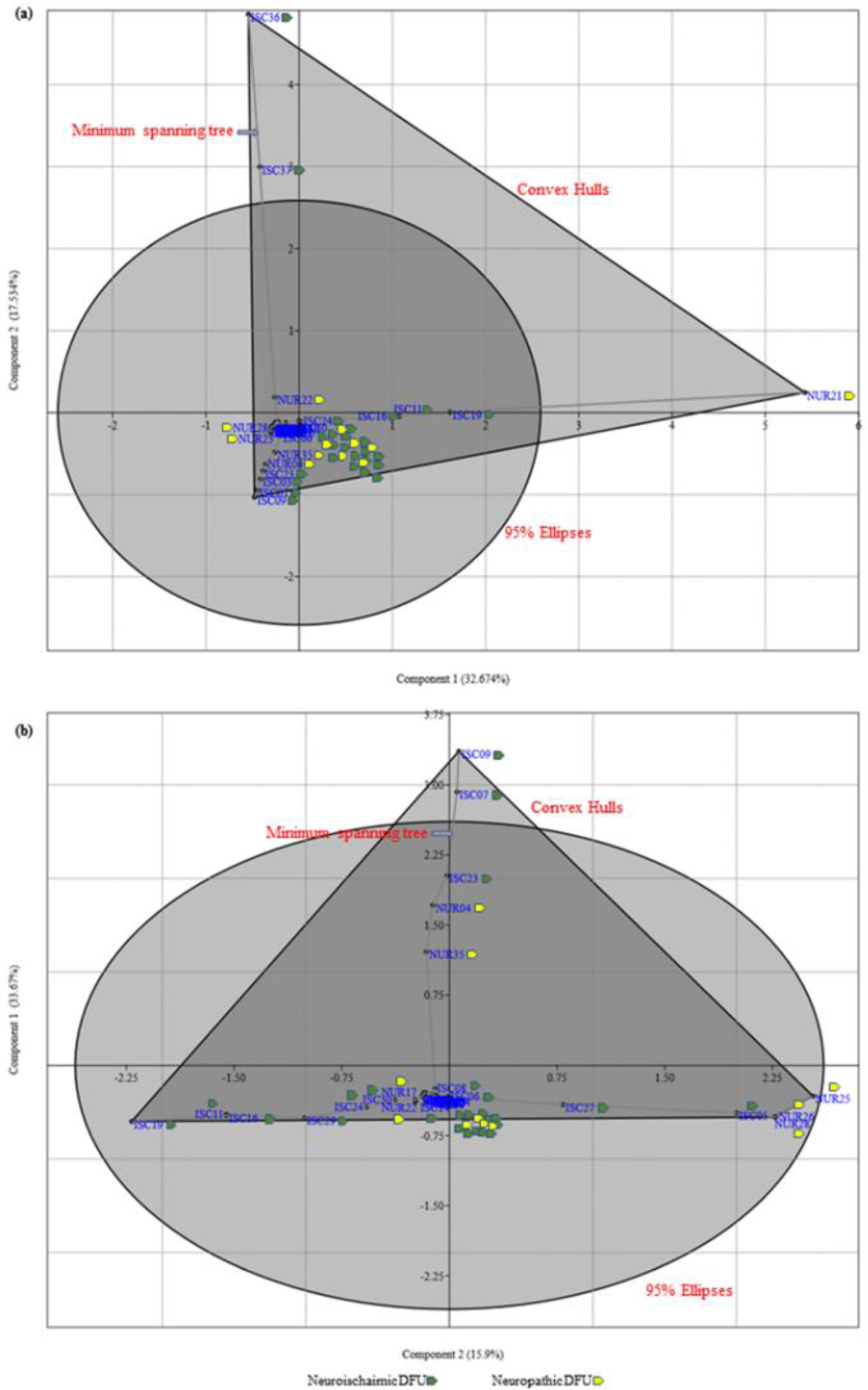
Principal component analysis (PCA) based on both qualitative and quantitative bacterial distribution in the DFU microbiome (a) and after removing the outliers (b).

## 4 Discussion

Diabetic foot ulcer (DFU) microbiome is still considered a pivotal point in wound infection. Indeed, the feet of healthy men versus diabetic men show a less presence of *Staphylococcus* germs, an increased presence of *S. aureus*, and more bacterial mixture (29). However, when weighed with the juxtaposed healthy skin, the DFU microbiome reveals fewer bacterial mixture but with more unscrupulous pathogens (30). Nevertheless, debatable outcomes have been recently reported with enhanced DFUs bacterial diversity and showing the inadequate link of well-known pathogens and specific different microbial patterns (31). Above that, DFUs has been linked to more diversified microbiomes (10, 32), having more anaerobic bacteria (23) when contrasted to other wounds. Surprisingly, in a recent study comparing the bacterial load of DFUs (n = 910), venous leg ulcers (n = 916), decubitus ulcers (n = 767), and chronic surgical wounds (n = 370), all lesions types show a similar degree of diversity and presence. Above that, the richness of pathogens in almost every wound was approximately the same [13] (33). This means those patient demographics and wound type had no impact on the chronic wound microbiome’s bacterial composition. Additional studies are required to confirm these results and demonstrate whether such wound types could be grouped into a single clinical management entity to improve wound management outcomes. Generally, we know that the ecology of the DFUs varies between the regions of the world: in western countries, gram-positive aerobic cocci remain the main germs, and in warmer climates, gram-negative bacilli are more predominant (29). Of course, this could affect the bacterial community structure of DFU microbiome. In the present study we investigated the DFU microbial load, microbial diversity, pathogenicity detected by 16S rRNA gene sequencing obtained from direct swab (DS) and cultured microbiomes (CM) against conventional microbiology culturing techniques routinely used in clinical setups. (In the same time the first consensus Saudi DFU microbiome). In addition, we compared between neuroischaimic and neuropathic DFU core microbiomes. We observed the presence of 26 species in the total DFU core microbiome (95% of reads of 38 individuals), utilizing both sampling methods. *Staphylococcus lugdunensis* was the most abundant species in the cultured DFU core microbiome method (18%). Additionally, the microbiome methods were able to present *Staphylococcus lugdunensis* as a new DFU causal agent since it represented the highest fraction of the core DFU microbiome in 5 cases (13%) in CM method and in 2 cases (5%) using DSM method. Furthermore, *S. lugdunensis* was part of the core microbiome in 32 cases (84%) for the CM method and 24 cases (63%) for DSM method. *S. lugdunensis* is a member of the coagulase negative staphylococci (CoNS) regularly inhabiting the human skin and mucosal membranes that can cause serious infections, with clinical characteristics that resemble those of *S. aureus*. Among the CoNS, only a few species are known to cause human disease, usually in the form of opportunistic infections only (34). Though, *S. lugdunensis* is an important exemption (35). *S. lugdunensis* is the causal agent of catheter-related bacteremia similar to other CoNS and it causes a variety of severe nosocomial and community-acquired infections, including native valve endocarditis, a devastating and potentially fatal disease that can affect previously healthy individuals. In addition, *S. lugdunensis* isolates are usually susceptible to multiple antimicrobial agents in comparison with other multiple-drug-resistant CoNS and *S. aureus* (36–38). It is worth mentioning that *S. lugdunensis* is a biosafety level two pathogen, and appropriate safety procedures should always be utilized while working with it (39). However, information about the clinical significance of *S. lugdunensis* is scarce. Moreover, *S. aureus* was the most abundant species in the direct swab DFU core microbiome method (21%). *S. aureus* is the leading common cause of skin and soft tissue infections. However, when it infiltrates blood after penetrating the skin, it can reach and infect bones and even heart valves (40). Nevertheless, *S. aureus* temporarily colonizing skin and mucosae in at least 25 to 30% of the population worldwide, without any clinical signs (41). However, in diabetic patients, *S. aureus* can change from an opportunistic colonizer to a pathogen involved in characteristic clinical manifestations, such as diabetic foot ulcers (42, 43). On the other hand, the unresolved OUTs that belong to *Staphylococcus sp.* was the least abundant component of the DFU cultured core microbiome (0.16%). While *Acinetobacter schindleri* was the least abundant component of the DFU core microbiome (0.014%) of the direct swab microbiome method. *A. schindleri* is a nonmotile, aerobic, Gram-negative bacterium (44). it is a ubiquitous opportunistic human pathogen affecting patients in intensive care units and immunocompromised patients and contributing to nosocomial infections and outbreaks worldwide because of its capability to survive on inert surfaces for a long duration (45–47). There is an underestimation of *A. schindleri* infections because of the indiscriminative nature of its phenotypic identification (48). Furthermore, *S. epidermidis* must not be disregarded as a potential DFU pathogen since it appeared as the highest fraction of the core DFU microbiome in 5 cases (13%) using the DSM in three cases (8%) using the CM method. *S. epidermidis* was part of the core microbiome in 21 cases (55%) for the CM method and 18 cases (47%) for the DSM method. Indeed, there is a great need to fully understand the pathogenic and protective roles of *S. epidermidis* before making any therapeutic decision involving it (49). It was found that *S. epidermidis* overexpansion and EcpA production are linked to aggravation of Netherton syndrome (NS), which is a skin ailment characterized by elevated levels of serine protease activity triggered by a *SPINK5* gene mutation (50). Moreover, *S. epidermidis* protease EcpA can be a deleterious component of the skin microbiome in atopic dermatitis. The overabundance of *S. epidermidis* found on some atopic patients can act similarly to *S. aureus* and damage the skin by expression of a cysteine protease (51). It was found that amongst *S. epidermidis* isolates, the disease-causing *isolates* embody a pathogenic sub-population that has attained genetic factors, horizontal gene transfer, and its related phenotypes that facilitates infection. Méric et al. identified 61 genes containing infection-associated genetic elements correlated with pathogenicity behaviors such as biofilm formation, cell toxicity, interleukin-8 production, and methicillin resistance (52). Furthermore, *Staphylococcus warneri* represented the highest fraction of the core DFU microbiome in 2 cases (5%) in the CM method, while it represented the highest fraction of the core DFU microbiome in one case (2.6%) using the DSM method. However, *S. warneri* was part of the core microbiome in 15 cases (39%) for the CM method and 12 cases (31.6%) for the DSM method. Coagulase-negative staphylococci, such as *S. warneri*, are skin commensals (53). However, there has been an increased acknowledgment of these organisms’ clinical significance in cultures. Clinically significant infection risk factors with *S. warneri* include immunosuppression, prosthetic devices, and intravascular catheters (54). Furthermore, among several detected *Corynebacterium species, C. striatum* and *C. amycolatum* are important DFU pathogens. *C. striatum* represented the highest fraction of the core DFU microbiome in one case (8%) in the CM method. Moreover, *C. striatum* was part of the core microbiome in 5 cases (13%) for the CM method and in 6 cases (15.8%) for the DSM method. Several 16S rRNA gene sequencing and shotgun metagenomics based studies showed that *Corynebacterium* was the second most abundant genera consistent with chronic wound microbiome and have a more significant role in wound healing than simple contamination from intact skin (33, 55, 56). In fact, it was found that *C. striatum*, typically considered commensals or contaminants, significantly impact both DFU wound severity and healing (56). Additionally, Patel et al., suggested that clinicians should consider *C. striatum* as a possible cause of osteomyelitis in diabetic patients, especially when patients fail to completely heal wounds in a timely manner that have previously and repeatedly displayed Diphtheroids from cultures (57). Furthermore, Hahn et al. showed that *C. striatum* is the most prevalent and abundant Corynebacterium spp. in DFU, detected in 28% of our specimens and is associated with infection and multi-drug resistance (58). Furthermore, *C. striatum* was presented as the causal agent of empyema and osteomyelitis in a patient with advanced rheumatoid arthritis (59). For, *C. amycolatum,* it appeared as the highest fraction of the core DFU microbiome in one case (8%) in CM method and in 2 cases (5%) for DSM method. It was part of the core microbiome in 10 cases (26%) for the CM method and in 14 cases (36.8%) for the DSM method. *C. amycolatum* was comparatively present in lesser abundances in DFU wounds (56). However, most recently Fernandez et al., reported two cases of mastitis caused by *C. pyruviciproducens* and *C. amycolatum* are described in not breastfeeding immunocompetent women and emphasizing on the importance to this genus by always ranking cultures, starting with a thorough sample collection up until a complete evaluation of lab results and clinical presentation (60). Moreover, one important pathogen is *Acinetobacter radioresistens* that requires close attention. It appeared as highest fraction of the core DFU microbiome in three cases (8%) in CM method. *A. radioresistens* was part of the core microbiome in 4 cases (10%) for the CM method. *A. radioresistens* is non-motile, non-sporulent, aerobic, Gram-negative rod-shaped bacterium, which is resistant to radiation, desiccation and carbapenem antibiotics. It is a constituent of the normal human skin microflora and an opportunistic pathogen in immunocompromised patients (61–64). To our knowledge, there are only four reports in the literature describing the isolation of *A. radioresistens* from human clinical specimens. Most recently, Wang et al., reported *A. radioresistens* infection with bacteremia and pneumonia. They emphasized on that *A. radioresistens* is the source of the Class D OXA-23 carbapenemase that can confer carbapenem resistance in *A. baumannii* and Thus, accurate identification of *A. radioresistens* is important for clinical management and to potentially prevent the spread of carbapenem resistance (65). Surprisingly, *Pseudomonas aeruginosa,* the well-established DFU pathogen, represented the highest fraction of the core DFU microbiome in only one case (DFU 31) (2.6%) in both microbiome methods. Indeed, *P. aeruginosa* is one of the major pathogens associated with the acute tissue damage in patients having DFU (66). Along with *S. aureus*, *P. aeruginosa* are the predominant Gram-positive and Gram-negative pathogens detected in DFIs (23, 67–69). As we stated before, conventional microbiological technique failed to compete with either tested molecular/microbiome methods in reflecting the real microbial situation inside the DFU in all 38 cases. On contrast with microbiome approaches, conventional microbiological technique was capable to detect only 1 or 2 pathogens, and in some cases no pathogen was distinguished. Oates et al., concluded that whereas chronic wounds commonly sheltered higher eubacterial diversity than healthy skin, the isolation of common pathogens was not associated with qualitatively discrete consortial patterns or altered diversity. The supported usage of both conventional microbiological techniques and molecular techniques, i.e., DGGE, for the microbial description of chronic wounds (31). Clinical laboratory investigations of chronic wound infections commonly depend on bacterial isolation by culture, which most efficiently detects dominant microbes that can grow on synthetic media. Whereas this is a valuable and well-established process for detecting numerous known pathogens associated with wound infections, it may underrate microbial diversity (70). Thus, reliance on molecular/genomics/next-generation sequences techniques will enable better resolution for DFU/chronic wound infections. A growing list of pathogens is becoming harder, and sometimes impossible, to treat as antibiotics become less effective. Antibiotics can be bought for human use without a prescription; sometimes, the emergence and spread of resistance are made worse. Similarly, in countries without standard treatment guidelines, antibiotics are often over-prescribed by health workers can exacerbate the burden of treating the diabetic infection that is one of the leading cause of limb loss among diabetic patients; so, having an accurate tool to target appropriate antibiotic therapy would a crucial if the most vital tool in this regards. Our current study shows that there is no big difference between the both used methods for wound culture but underlined the importance of culturing wound as mandatory wound assessment for appropriate antibiotic therapy, therefore targeting the main involved pathogens that usually impede wound healing process. Generally, the NURDFU group showed higher values for the tested diversity indices than the ISCDFU group. The number of observed taxa ranged between 19-66 with a mean of 34.5769 taxa with a sum of 899 for the ISCDFU group. Neither Cluster analysis UPGMA (Correlation, elucidation or Bary-Curtis criteria) (Figure 12) or Principal component analysis (PCA) (Figure 13a and b), based on both qualitative and quantitative bacterial distribution, showed no clear clustering or pattern amongst the two types of ulcers, namely, NURDFU and ISCDFU. Moreover, Generally, the investigated clinical factors distributed arbitrarily among the observed 13 clusters. However, a larger sized investigation is still required to validate our finding. Wound infection is a key factor as well as vasculopathy and Neuropathy in the prevention of limb loss, so swab wound culture at the patient side has to be a standard in monitoring wound care and guiding treatment. “No need to kill a mosquito with a bomb” appropriate swab culture is a cost-effective and adequate tool for diabetic wound management to avoid amputation. Diabetic foot ulcer (DFU) microbiome remains a key point in wound infections; using appropriate would culture tool will not save time but also reduce the diabetic foot burden as well as saving time, money and prevent lower limb amputation among the Saudi diabetic population where one out of fore is diabetic.

## 5 Conclusions

DFIs are considered a significant public health problem in wound management. Prompt assessment and adequate treatment are a cornerstone in preventing limb loss among the diabetic population. This study underlined the differences between traditional microbiological analysis, direct swab (DSM), and cultured (CM) to assess the DFU microbiome for an appropriate bacterial burden analysis to direct treatment and prevent infection. Our results showed that cultured microbiome is superior to both the traditional method and direct swab microbiome. Generally, the Neuroischaimic group showed higher values for the tested diversity indices than the Neuroischaimic group. Neither Cluster Analysis nor Principal Component Analysis (PCA) showed apparent clustering or pattern amongst the two types of ulcers.

Nevertheless, new data derived from molecular tools suggest that chronic wounds contain consortia of similar pathogens coexisting as combinations of highly structured communities. This current study provides a simple, affordable, and cost-effective tool, direct wound sampling, to uncover the specific roles and treatment of these pathogens in chronic wounds. Moreover, further studies using similar methods of microorganism identification and well-defined populations with DFIs together with standardized sampling methods will enable a better understanding of the bacterial diversity, will provide new insights to redirect treatments, and might improve clinical outcomes in the future.

## Supplemental Figures

**Supplementary Figure 1**: Summarized Research methodology

## Supplementary Tables

**Supplementary table 1** clinical and demographic information of the study subjects

**Supplementary Table 2** Distribution of the observed DFU microbiome species (OTUs) among the total Cultured Microbiome and Direct Swap Microbiome methods.

**Supplementary Table 3** The exclusive 40 species detected in Cultured Microbiome only

**Supplementary Table 4** The exclusive 21 species detected in Direct swap Microbiome only.

**Supplementary Table 5** Diversity indices for the 26 subjects with neuroischaimic DFU (NISCDFU).

**Supplementary Table 6** Diversity indices for the 12 subjects with neuropathic DFU (NUR).

## * List of abbreviations

DFU: Diabetic foot ulcer
PEDIS: Perfusion, Extent, Depth, Infection, and Sensation
DSM: direct swab microbiome method
CM: cultured microbiome method
ABPI: Ankle-brachial pressure index
ISCDFU: neuroischaimic DFU
NURDFU: neuropathic DFU
NGS: Next-generation sequencing techniques
16S rRNA: 16S ribosomal RNA gene
Mb: Megabase pairs
BLASTn: Basic Local Alignment Search Tool nucleotide
OTUs: operational taxonomic units
UPGMA: unweighted pair group method with arithmetic mean
PCA: Principal component analysis

## Declarations

### **\*** Ethics approval and consent to participate

The institutional review board approved this study at King Saud University, College of Medicine Riyadh, Kingdom of Saudi Arabia (IRB-E-16-1839). The subjects provided written informed consent for participating in this study.

### **\*** Consent to publish

All authors have consented for the publication of this manuscript.

### * Availability of data and materials

All sequence data were submitted to NCBI with an SRA accession: SUB4604842 and BioSample accessions numbers from SAMN10174890 to SAMN10174927 under the BioProject PRJNA494939

### **\*** Authors’ contributions

**AF** Admistratioanal supervision of the research and preparing the final approval of the version to be published.

**ATMS** Involved in study conception and design, data analysis, and interpretation and involved in drafting the manuscript or revising it critically for important intellectual content. Preparing the final approval of the version to be published.

**AY** Involved in data and statistical analysis.

**BM** Performed the DNA sequencing and Involved in drafting the manuscript.

**HT** Involved in study conception and design and drafted the manuscript or revised it critically for important intellectual content. She was involved in preparing the final approval of the version to be published.

**KA** Involved in study conception and design. Preparing the final approval of the version to be published.

**MN** Providing research fund for the research and final approval of the version to be published.

**UR** Involved in the acquisition of data or analysis and interpretation of data in drafting the manuscript.

## Acknowledgment

The authors want to thank the members of the University Diabetes Center at King Saud University for their help throughout the study. Special thanks for Prof. Khalid Al-Rubeaan for his great help in this work. We thanks to Mrs. Hessa Al-Brahim and Mr. Mohthash Musambil for their assist in laboratory work. Furthermore, we want to thank King Saud University Medical City, King Saud University, Riyadh, Saudi Arabia, for assisting in conducting the research. Besides, we want to acknowledge that NGS experiments and analysis were supported by the Saudi Human Genome Program (SHGP) at KACST and KFSHRC.

